# A meta-analytic approach to mapping co-occurrent grey matter volume increases and decreases in psychiatric disorders

**DOI:** 10.1101/2020.05.18.101436

**Authors:** Lorenzo Mancuso, Alex Fornito, Tommaso Costa, Linda Ficco, Donato Liloia, Jordi Manuello, Sergio Duca, Franco Cauda

## Abstract

Numerous studies have investigated gray matter (GM) volume changes in diverse patient groups. Reports of disorder-related GM reductions are common in such work, but many studies also report evidence for GM volume increases in patients. It is unclear whether these GM increases and decreases independent or related in some way. Here, we address this question using a novel meta-analytic network mapping approach. We used a coordinate-based meta-analysis of 64 voxel-based morphometry studies of psychiatric disorders to calculate the probability of finding a GM increase or decrease in one region given an observed change in the opposite direction in another region. Estimating this co-occurrence probability for every pair of brain regions allowed us to build a network of concurrent GM changes of opposing polarity. Our analysis revealed that disorder-related GM increases and decreases are not independent; instead, a GM change in one area is often statistically related to a change of opposite polarity in other areas, highlighting distributed yet coordinated changes in GM volume as a function of brain pathology. Most regions showing GM changes linked to an opposite change in a distal area were located in salience, executive-control and default mode networks, as well as the thalamus and basal ganglia. Moreover, pairs of regions showing coupled changes of opposite polarity were more likely to belong to different canonical networks than to the same one. Our results suggest that regional GM alterations in psychiatric disorders are often accompanied by opposing changes in distal regions that belong to distinct functional networks.

## Introduction

A large body of neuroimaging studies has investigated how diverse diseases are associated with altered brain structure, most commonly quantified through measures of regional grey matter (GM) volume. The vast majority of studies have focused on mapping localized changes using mass univariate approaches such as voxel-based morphometry (Ashburner and Friston, 2000), but analyses of covariations in regional volume changes are also thought to reveal pathological mechanisms and to reflect the distributed and interconnected nature of the brain (Evans, 2013). By far, most work in this area has focused on understanding GM volume reductions in clinical disease. Not only have several meta-analyses of different diseases shown that such reductions are common (Fornito *et al*., 2009; Bora *et al*., 2010, 2011, 2012*a, b*; Fusar-Poli *et al*., 2011; Hallahan *et al*., 2011; Linkersdörfer *et al*., 2012; Du *et al*., 2012; Li *et al*., 2014, 2018; Stoodley, 2014; Cauda *et al*., 2014; Foster *et al*., 2015; Lin *et al*., 2016; Wise *et al*., 2017; Wu *et al*., 2018), and other work suggests that anatomically distributed yet coordinated GM reductions are tied to the underlying connectivity between regions (Seeley *et al*., 2009; Raj *et al*., 2012; Zhou *et al*., 2012; Crossley *et al*., 2014; Iturria-Medina *et al*., 2014; Zeighami *et al*., 2015, Cauda *et al*., 2018*a*; Manuello *et al*., 2018; Yau *et al*., 2018; Zheng *et al*., 2019). In contrast, GM increases are less commonly considered in clinical neuroimaging studies (Cauda *et al*., 2011, 2017, 2018*b*; Tatu *et al*., 2018, Cauda *et al*., 2019*b*; Ding *et al*., 2019; Lu *et al*., 2019), potentially because they might be a rarer consequence of disease and because they can be difficult to explain in the context of pathology. Indeed, while a morphometric decrease can be easily interpreted as a sign of neurodegeneration or neurodevelopmental hyperpruning, the interpretation of a disorder-related GM increase is less clear.

Candidate mechanisms for increased GM include modifications to neuronal tissue, such as neurogenesis (Eriksson *et al*., 1998), synaptogenesis (Sarrazin *et al*., 2019) or changes in somal size and density, in addition to changes in glia (Rocha *et al*., 1998) or neurovasculature (Zatorre *et al*., 2012). There could be diverse causes of such modifications, which may be related to inflammatory processes (Poletti *et al*., 2019) that might, for instance, induce astrocytic hypertrophy (Li *et al*., 2019), or to activity-driven volumetric increases similar to those observed during learning (Zatorre *et al*., 2012). Medications may also an hypertrophic effect (Torres *et al*., 2013). Otherwise, a GM increase in patients compared to controls could derive from a pathologic hypopruning that could characterize neurodevelopmental diseases such as schizophrenia (Keshavan *et al*., 1994*a*). More trivial reasons, such as case-control differences in hydration, head motion, and various other metabolic and physical confounds may also play a role (Weinberger and Radulescu, 2016).

Some authors hypothesised that GM increases could emerge as a compensatory response to localised damage elsewhere. For instance, Janson and colleagues (Janson *et al*., 1991) observed neuronal hypertrophy in regions connected to a mesencephalic lesion, which they interpreted as a compensation to the injury. Stevens and colleagues (Stevens, 1992) proposed that hippocampal lesions might produce axonal sprouting and synapse proliferation in deafferented regions, and that these changes should have a compensatory function if the rewiring is adaptive. However, they could also lead to a further impairment if the adaptation is suboptimal. In major depression, cortical thickness studies suggest that an early phase of the disease could be characterized by an increased thickness in some regions, in which provisional compensatory mechanisms take place to overcome the deficits induced by the damage in others (Qiu *et al*., 2014; Li *et al*., 2019). Indirect evidence for the existence compensatory changes in GM volume comes from a meta-analysis (Cauda *et al*., 2014) showing that, in people with autism, a GM decrease in one area is associated with an increase of volume and/or DTI-related measures of connectivity in the white matter tracts connecting the affected area. In functional neuroimaging, several studies suggest the presence of compensatory activations during both healthy ageing and in diverse diseases (eg. Dolcos *et al*., 2002; Tan *et al*., 2006; Crossley *et al*., 2016). For instance, after traumatic brain injury, increased functional connectivity of default mode network (DMN), salience network (SN) and executive control network (ECN) has been observed (Hillary *et al*., 2014). These findings align with the view that the interconnected architecture of the brain means that pathology is seldom defined to a single locus, and may induce distributed responses that are both adaptive and/or maladaptive (Fornito *et al*., 2015).

If compensatory changes in GM volume do occur in the diseased brain, or if disorder-related GM increases and decreases are more broadly coordinated in some way, then we should expect that these changes should be statistically associated across different disorders. To test this hypothesis, we extended a coordinate-based meta-analytic methodology developed by our group (Cauda *et al*., 2018*b*; Manuello *et al*., 2018; Tatu *et al*., 2018) to calculate the probability of co-occurence between GM increases and decreases in VBM studies of psychiatric disease. This probability quantifies the likelihood that a GM increase or decrease in one brain area co-occurs with a GM change of opposite polarity in another area. We adopted a transdiagnostic approach, considering all the brain disorders that would show an association between GM decreases and increases, to identify disease-invariant co-alteration mechanisms of brain pathology (Buckholtz and Meyer-Lindenberg, 2012, Cauda *et al*., 2018*b*). Thus, we aimed to obtain a network of GM co-alterations of opposing polarity (COA-O) that identifies pairs of regions for which there is a statistical dependence between these two opposite forms of GM change. Given prior work indicating that networks of co-alterations (i.e., GM changes of the same polarity) are related to normative connectivity patterns (Yates, 2012, Cauda *et al*., 2018*b*; Raj and Powell, 2018), we further hypothesised that the COA-O network would show a significant correlation with functional connectivity (FC) networks in healthy individuals.

## Material and Methods

### Data collection

We adopted the Cochrane Collaboration definition of meta-analysis (Green *et al*., 2008) and the “PRISMA statement” international guidelines for the selection of studies (Liberati *et al*., 2009; Moher *et al*., 2009). Coordinates of statistically significant GM changes were obtained from the BrainMap database (http://brainmap.org/) (Fox and Lancaster, 2002; Fox *et al*., 2005; Laird *et al*., 2005; Vanasse *et al*., 2018).

Two queries were conducted on the VBM BrainMap section (November 2019) to retrieve data on morphometric decreases and increases, respectively:

1. [Experiments Context is Disease] AND [Experiment Contrast is Gray Matter] AND [Experiments Observed Changes is Controls>Patients];
2. [Experiments Context is Disease] AND [Experiment Contrast is Gray Matter] AND [Experiments Observed Changes is Patients>Controls].

We obtained 1001 studies reporting GM decreases and 382 studies reporting GM increases. All the selected experiments with a sample size smaller than 8 participants were eliminated, in line with Scarpazza et al. (2015), who showed that VBM experiments with more than 8 subjects should not be biased by an increased false positive rate. Then we coded the experiments according to the ICD-10 classification (World Health Organisation, 1992), to exclude the non-neurologic and non-psychiatric diseases from the database. In order to quantify the co-occurrence of decreases and increases, we further selected only those couples of experiments (i.e. set of foci resulting by a given statistical comparison) that reported opposing changes between the same groups of patients and healthy controls. Such selection resulted in 170 experiments (85 decreases exp., 85 increases exp.), further reduced to 128 after the exclusion of neurological diseases. See Table 1 and 2 for a list of the experiments and disorders considered.

**Table 1:** summary of the pathologies obtained by the first selection of studies. Experiments about psychiatric disorders were included in the meta-analysis, while those about neurological disorders were excluded.

| **included** | **Diagnosis** | **ICD-10 code** | **Number of experiments for condition** |
| --- | --- | --- | --- |
| yes | Schizophrenia | F20 | 25 |
| yes | Pervasive developmental disorders, autism | F84 | 13 |
| yes | Bipolar disorder | F31 | 7 |
| yes | Major depressive disorder | F31-33 | 7 |
| yes | Obsessive-compulsive disorder | F42 | 6 |
| yes | Attention deficit/hyperactivity disorder | F90 | 2 |
| yes | Specific developmental disorders of speech and language | F80 | 2 |
| yes | Other disorders of psychological development, Tourette syndrome | F95 | 1 |
| yes | Other psychotic disorder not due to a substance or known physiological condition | F28 | 1 |
| no | Epilepsy and Recurrent Seizures TLE, JME | G40 | 13 |
| no | Other extrapyramidal and movement disorders | G25 | 4 |
| no | Dystonia | G24 | 2 |
| no | Parkinson's Disease PD, ET | G20 | 2 |

**Table 2:** list of experiments obtained by the first selection of studies.. Each entry of the list represents an experiment of decrease and one of increase. Experiments about psychiatric disorders were included in the meta-analysis, while those about neurological disorders were excluded.

| **included** | **author** | **# subjects** | **ICD-10 code** | **Diagnosis** |
| --- | --- | --- | --- | --- |
| yes | Abell F, 1999 | 15 | F84 | Pervasive Developmental Disorders PDD, autism, ASD |
| yes | Adler CM, 2005 | 27 | F31 | Bipolar Disorder BD, BPD |
| yes | Antonova E, 2005 | 40 | F20 | Schizophrenia SZ |
| yes | Arnone D, 2009 | 25 | F32-F33 | Major Depressive Disorder |
| yes | Arnone D, 2013 | 39 | F32-F33 | Major Depressive Disorder |
| yes | Bassitt DP, 2007 | 30 | F20 | Schizophrenia SZ |
| yes | Brieber S, 2007 | 15 | F84 | Pervasive Developmental Disorders PDD, autism, ASD |
| yes | Brieber S, 2007 | 15 | F90 | Attention Deficit/Hyperactivity Disorder |
| yes | Cheng Y, 2011 | 25 | F84 | Pervasive Developmental Disorders PDD, autism, ASD |
| yes | Cheng Y, 2011 | 11 | F84 | Pervasive Developmental Disorders PDD, autism, ASD |
| yes | Cheng Y, 2011 | 12 | F84 | Pervasive Developmental Disorders PDD, autism, ASD |
| yes | Cheng Y, 2011 | 12 | F84 | Pervasive Developmental Disorders PDD, autism, ASD |
| yes | Cui L, 2011 | 23 | F20 | Schizophrenia SZ |
| yes | Cui L, 2011 | 24 | F31 | Bipolar Disorder BD, BPD |
| yes | Deng MY, 2009 | 10 | F20 | Schizophrenia SZ |
| yes | Deng MY, 2009 | 10 | F20 | Schizophrenia SZ |
| yes | Ecker C, 2010 | 22 | F84 | Pervasive Developmental Disorders PDD, autism, ASD |
| yes | Ecker C, 2012 | 89 | F84 | Pervasive Developmental Disorders PDD, autism, ASD |
| yes | Gilbert AR, 2008 | 20 | F42 | Obsessive Compulsive Disorder OCD |
| yes | Giuliani NR, 2005 | 34 | F20 | Schizophrenia SZ |
| yes | Gong Q, 2011 | 23 | F32-F33 | Major Depressive Disorder |
| yes | Gong Q, 2011 | 23 | F32-F33 | Major Depressive Disorder |
| yes | Haldane M, 2008 | 44 | F31 | Bipolar Disorder BD, BPD |
| yes | Ha TH, 2004 | 35 | F20 | Schizophrenia SZ |
| yes | Honea RA, 2008 | 169 | F20 | Schizophrenia SZ |
| yes | Hulshoff Pol HE, 2001 | 158 | F20 | Schizophrenia SZ |
| yes | Hwang J, 2010 | 26 | F32-F33 | Major Depressive Disorder |
| yes | Hyde KL, 2010 | 13 | F84 | Pervasive Developmental Disorders PDD, autism, ASD |
| yes | Kasperek T, 2010 | 49 | F20 | Schizophrenia SZ |
| yes | Kawasaki Y, 2004 | 25 | F20 | Schizophrenia SZ |
| yes | Ke X, 2008 | 15 | F84 | Pervasive Developmental Disorders PDD, autism, ASD |
| yes | Ladouceur CD, 2008 | 20 | F31 | Bipolar Disorder BD, BPD |
| yes | Leung KK, 2009 | 17 | F32-F33 | Major Depressive Disorder |
| yes | Lu C, 2010 | 12 | F80 | Specific Developmental Disorders of Speech and Language |
| yes | Ludolph AG, 2006 | 14 | F95 | Other Disorders of Psychological Development. Tourette |
| yes | Marcelis M, 2003 | 27 | F28 | Other psychotic disorder not due to a substance or known physiological condition |
| yes | Mengotti P, 2011 | 20 | F84 | Pervasive Developmental Disorders PDD, autism, ASD |
| yes | Molina V, 2011 | 24 | F20 | Schizophrenia SZ |
| yes | Molina V, 2011 | 30 | F20 | Schizophrenia SZ |
| yes | O'Daly O, 2007 | 28 | F20 | Schizophrenia SZ |
| yes | Price G, 2010 | 47 | F20 | Schizophrenia SZ |
| yes | Pujol J, 2004 | 72 | F42 | Obsessive Compulsive Disorder OCD |
| yes | Salgado-Pineda P, 2003 | 13 | F20 | Schizophrenia SZ |
| yes | Salmond CH, 2007 | 9 | F84 | Pervasive Developmental Disorders PDD, autism, ASD |
| yes | Saricicek A, 2015 | 28 | F31 | Bipolar Disorder BD, BPD |
| yes | Scheuerecker J, 2010 | 13 | F32-F33 | Major Depressive Disorder |
| yes | Schiffer B, 2013 | 25 | F20 | Schizophrenia SZ |
| yes | Shapleske J, 2002 | 31 | F20 | Schizophrenia SZ |
| yes | Shapleske J, 2002 | 32 | F20 | Schizophrenia SZ |
| yes | SmesnyS,2010 | 13 | F20 | Schizophrenia SZ |
| yes | Smesny S, 2010 | 11 | F20 | Schizophrenia SZ |
| yes | Suzuki M, 2002 | 42 | F20 | Schizophrenia SZ |
| yes | Szeszko PR, 2008 | 26 | F42 | Obsessive Compulsive Disorder OCD |
| yes | Tang LR, 2014 | 27 | F31 | Bipolar Disorder BD, BPD |
| yes | Toal F, 2010 | 26 | F84 | Pervasive Developmental Disorders PDD, autism, ASD |
| yes | Valente AA Jr, 2005 | 15 | F42 | Obsessive Compulsive Disorder OCD |
| yes | Valente AA Jr, 2005 | 15 | F42 | Obsessive Compulsive Disorder OCD |
| yes | Wang J, 2007 | 12 | F90 | Attention Deficit/Hyperactivity Disorder |
| yes | Watkins KE, 2002 | 10 | F80 | Specific Developmental Disorders of Speech and Language |
| yes | Watson DR, 2012 | 25 | F20 | Schizophrenia SZ |
| yes | Watson DR, 2012 | 24 | F31 | Bipolar Disorder BD, BPD |
| yes | Whitford TJ, 2006 | 41 | F20 | Schizophrenia SZ |
| yes | Wilke M, 2001 | 48 | F20 | Schizophrenia SZ |
| yes | Yoo SY, 2008 | 47 | F42 | Obsessive Compulsive Disorder OCD |
| no | Celle S, 2010 | 17 | G25 | Other extrapyramidal and movement disorders |
| no | Chan CH, 2006 | 13 | G40 | Epilepsy and Recurrent Seizures TLE, JME |
| no | De Araujo-Filho GM, | 16 | G40 | Epilepsy and Recurrent Seizures TLE, JME |
|  | 2009 |  |  |  |
| no | Etgen T, 2005 | 28 | G25 | Other extrapyramidal and movement disorders |
| no | Granert O, 2011 | 11 | G24 | Dystonia |
| no | Keller SS, 2002 | 40 | G40 | Epilepsy and Recurrent Seizures TLE, JME |
| no | Keller SS, 2002 | 36 | G40 | Epilepsy and Recurrent Seizures TLE, JME |
| no | Keller SS, 2002 | 58 | G40 | Epilepsy and Recurrent Seizures TLE, JME |
| no | Keller SS, 2002 | 58 | G40 | Epilepsy and Recurrent Seizures TLE, JME |
| no | Keller SS, 2002 | 58 | G40 | Epilepsy and Recurrent Seizures TLE, JME |
| no | Kim JH, 2007 | 25 | G40 | Epilepsy and Recurrent Seizures TLE, JME |
| no | Lin CH, 2013 | 10 | G20 | Parkinson's Disease PD, ET |
| no | Lin CH, 2013 | 10 | G20 | Parkinson's Disease PD, ET |
| no | Lin CH, 2013 | 10 | G25 | Other extrapyramidal and movement disorders |
| no | Lin CH, 2013 | 10 | G25 | Other extrapyramidal and movement disorders |
| no | Lin K, 2009 | 30 | G40 | Epilepsy and Recurrent Seizures TLE, JME |
| no | Lin K, 2009 | 19 | G40 | Epilepsy and Recurrent Seizures TLE, JME |
| no | Lin K, 2009 | 30 | G40 | Epilepsy and Recurrent Seizures TLE, JME |
| no | Obermann M, 2007 | 9 | G24 | Dystonia |
| no | Riederer F, 2008 | 12 | G40 | Epilepsy and Recurrent Seizures TLE, JME |
| no | Riederer F, 2008 | 12 | G40 | Epilepsy and Recurrent Seizures TLE, JME |

To obtain two control datasets of common change (i.e., datasets in which only GM decreases or only GM increases were reported), we selected from the total pool of 1382 experiments, those considering the 9 conditions we focus on, obtaining 280 experiments for GM decreases and 115 experiments for GM increases (Note: this selection did not require the constraint that increases and decreases be reported in the same study; see Fig. S1 for the PRISMA flow chart and Tables S1 and S2 for the lists of experiments included in the control analyses). We will refer to those two control network of decrease-only and increase-only as COA-D and COA-I, respectively.

### Quantifying co-alteration probabilities

Our method is based on the Anatomical Likelihood Estimation (ALE) (Eickhoff *et al*., 2009, 2012; Turkeltaub *et al*., 2012). The ALE is a coordinate-based meta-analytic technique that aims to produce a map as the union of a set of modelled alteration (MA) maps, each one representing one statistical comparison (i.e. experiment) included in the study. For each experiment, its MA map is produced creating a 3-D Gaussian distribution of probability around each reported focus of alteration (Eickhoff *et al*., 2009).Their union produce a (unthresholded) ALE map, which represent the statistical distribution of the alterations across the experiments.

Our aim is to quantify the degree to which a GM change in one brain region is associated with a change of opposite polarity in another area. We do this for all possible pairs of brain regions, thus building a meta-analytic COA-O network. To build the network we need a unique set of nodes that can represent both the loci of both decreases and increases, and a set of maps that describe the alterations reported by the literature. To obtain the alteration maps from the foci retrieved from the BrainMap database, we computed a MA map for each experiment. To obtain the set of nodes, two unthresholded ALE maps were produced with GingerALE (http://www.brainmap.org/ale/) by merging the GM increase and GM decrease MA maps in the main dataset. When merging, we took the maximum value of the two maps for each voxel (Fig. 1 A). The merged map represents a common spatial distribution of the GM changes in our database. This map was then fed to a peak detection algorithm to identify the coordinates of alteration (Fig. 1 B). The nodes for the COA-D and COA-I were obtained using the same algorithm on the two ALE maps of the two control datasets, producing 233 nodes for the GM-decrease network and 269 nodes for the GM-increase network. Defining network nodes in this data-driven way allows us to more accurately sample the spatial locations of actual GM changes and, critically, to create equally-sized ROIs. In fact, each ROI was considered as altered in a given experiment if the 20% of its voxels were included in the corresponding MA map, thresholded at *p* <0.05(Fig 1 C). Since the probability indexes used in the analyses requires binary data of alteration (i.e. a node can be only altered or not, see below), this filtering step was necessary to avoid false positive alterations (i.e. labelling a node as altered if it contains only the periphery of a probability distribution). Although the 20% threshold is arbitrary, it has been previously shown that other thresholds do not change radically the results (Mancuso *et al*., 2019). Therefore, small ROIs are more likely to reach such threshold of alteration, thus having equally sized nodes would avoid such bias. However, to prove that our results would hold to different methodological choices, we also replicated the network using a pre-defined parcellation based on the Brainnetome Atlas (Fan *et al*., 2016).

**Figure 1:**
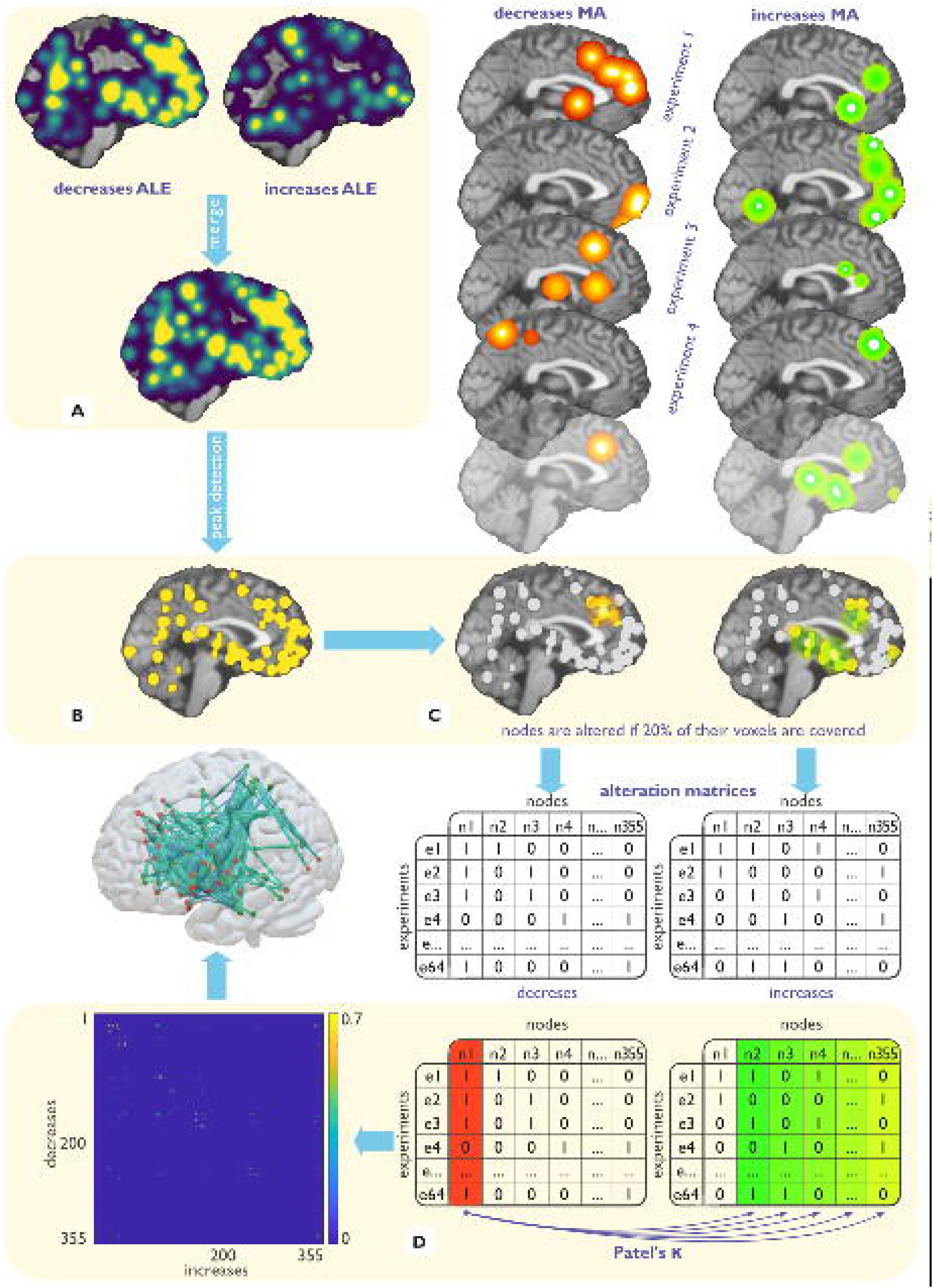
illustration of the methods for the calculation of the decrease-increase association matrix. **A**: the untresholded ALE maps of the decreases and the increases databases are merged. **B**: such map is fed to a peak detection algorithm to obtain the nodes of the network; **C**: each node is considered altered in each experiment if the 20% of its voxels are covered by a significant MA map voxel of that experiment. This operation produce a vector of 1 and 0 for each node that describes if that node is altered or not in each experiment, creating two alteration matrices, one for the decreases, one for the increases. **D**: the Patel’s κ is calculated between each node vector of the decreases alteration matrix and each node vector of the increases alteration matrix. This produces a matrix of co-alteration between the decreases and the increases, which gives us the intensity of the edges of the network.

This procedure was repeated on each dataset, resulting in 4 node x experiment matrices of GM changes (i.e., two paired 255×64 matrices of decrease-only and increase only for the main analysis, and two 233×280 and 269×115 matrices for the two decrease-only and increase-only control networks). The probability of observing a co-occurrent change in each pair of nodes was estimated using Patel’s κ (Patel *et al*., 2006). Table 3 illustrates the possible combinations of two given nodes *a* and *b*, along with their marginal probabilities.

**Table 3:** alteration states and marginal probability.

|  |  |  |  |  |
| --- | --- | --- | --- | --- |
| Node a | Node b |  |  |  |
|  |  | Altered | Non-altered |  |
| | Altered | $\vartheta_1$ | $\vartheta_3$ | $\vartheta_1 + \vartheta_3$ |
| | Non-altered | $\vartheta_2$ | $\vartheta_4$ | $\vartheta_2 + \vartheta_4$ |
| | | $\vartheta_1 + \vartheta_2$ | $\vartheta_3 + \vartheta_4$ | 1 |

Those marginal probabilities are essential for the Patel’s κ, which calculates the probability that the two nodes show co-occurring GM changes relative to the probability that they are altered independently, as

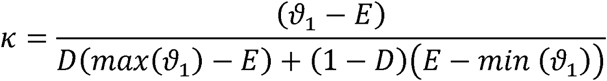

where

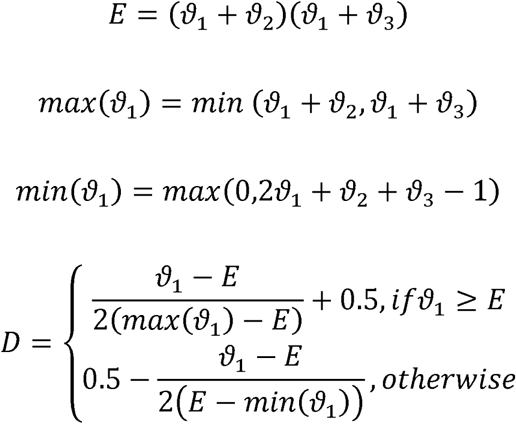

The numerator in the fraction is the difference between the likelihood that *a* and *b* are altered together and the expected likelihood *E* that *a* and *b* are co-altered under independence; *E* is the prior information of our Bayesian framework that, in a frequentist paradigm, would be disregarded or treated as not fixed by the data (Patel *et al*., 2006). The denominator is a weighted normalizing constant to restrict the Patel’s κ to the range [–1, 1]. The statistical significance (*p* = 0.01) is evaluated through a Monte Carlo simulation that calculates an estimate of *p*(*k*|*z*) by sampling a Dirichlet distribution and determining the proportion of the samples in which *k* > *e*, where *e* is the threshold of statistical significance. The resulting co-alteration matrix (be it COA-O, COA-I, COA-D) comprises values that are proportional to the statistical relationship between the alterations of the considered brain areas.

Crucially, while the co-alteration control matrix was obtained calculating κ between the 1,2,…N nodes within their respective nodes x experiments matrices of alteration (233×280 for the decrease condition, resulting in a 233×233 COA-D matrix, and p<0.05 for the increase condition, resulting in a 269×269 COA-I matrix), the COA-O matrix was produced calculating the κ between nodes 1,2,…355 of the 355×64 decreases matrix and the 1,2,…355 nodes of the 355×64 increases matrix (Fig. 1 D). It must be stressed that the number of nodes that are effectively connected in the final network is less than the 355 found with the peak detection algorithm, as some nodes have no significant edges after the statistical thresholding.

For each edge with a significant κ of the COA-O network, we also calculated Patel’s τ (Patel *et al*., 2006), which evaluates the asymmetries in conditional probabilities of a given pair of nodes as

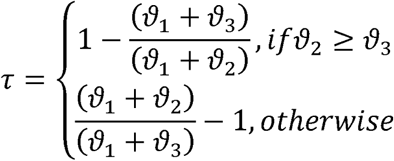

A positive value means that node *a* is ascendant to (i.e., has an influence on) node *b*. In practice, it means that when *a* shows an alteration, *b* usually do not; however, when *b* is altered, *a* tend to be altered as well. Therefore, if one of the two nodes is influencing the other, the direction is more likely to go from *a* to *b* than the other way around. In our implementation (Fig. 1 D), a positive value means a possible influence of a GM decrease on an increase.

### Resting state functional connectivity

Functional data were retrieved from the Cambridge dataset of the Functional Connectome Project (Biswal *et al*., 2010). The sample comprises 198 subjects (75M-123F, 18-30 years old). Each scan consists of 119 time points with TR=3. These data were processed with DPABI 3.1, DPARSFa 4.4 (http://rfmri.org/DPARSF) (Yan *et al*., 2016). The preprocessing steps were i) slice timing correction; ii) realignment; iii) regression of motion parameters using the Friston-24 model and of white matter and cerebrospinal fluid signals using *a priori* masks; iv) spatial normalization to standard SPM EPI template; v) smoothing with a 4 mm FWHM kernel; vi) scrubbing as in Power et al. (Power *et al*., 2012).

Then we extracted the functional signal of the 355 ROIs of the COA-O network to calculate the individual matrix of time series correlations, which were then normalized with the Fisher transformation and averaged into a group FC matrix.

To further investigate the relationship between the COA-O network and normative FC, each node was assigned to a resting state network (RSN) of the Yeo7 parcellation (Yeo *et al*., 2011), plus the cerebellum and a thalamus/basal ganglia masks (Thal/BG) created on the basis of the Brainnettome Atlas.

### Fail-safe and dummy-pairs analyses

We might hypothesize that the dataset of the experiments included in the COA-O analysis could be just a subsample of all the possible evidences about GM changes of opposite polarity, thus biasing our results. To assess their robustness, we implemented a modified version of the Fail-safe technique (Acar *et al*., 2018). The ratio behind that is that we cannot represent the COA-O network as if we gathered all the possible experiments on the matter, but we still can observe what would happen if our sample was much larger. Therefore, we generated 220 couples of random MAs using the fail-safe R script (https://github.com/NeuroStat/GenerateNull). Such MAs couples were progressively introduced in the database 10 at a time, recalculating the network after each injection. Then, the values of the edges of each new network were correlated with those of the original COA-O network. Since it is possible that any injection step could improve the correlation by mere chance, the method can produce random fluctuations in the correlation values, so the procedure was repeated 30 times. The more the network would hold to the injection of noise, the more it could be assumed that our network is an accurate representation of the COA-O phenomenon, even if our dataset was a subsample of a much larger set of experiments.

To calculate the co-alteration matrix, it was necessary to couple an experiment of decreases to the matching experiment of increases obtained on the same subjects. This forced us to discard all the papers that do not present the opposing statistical contrast between patients and healthy controls group. In general, this meant that we discarded many studies that only reported GM decreases in patients and provided no information about GM increases. There are two possible explanations as to why a study might only report GM decreases. One is that the researchers never tested for GM increases, despite the fact that they actually exist in the patient group. In this case, the data are incomplete and so it is necessary to discard these studies. The second reason is that the researchers ran the contrast, found no evidence of GM increases, and then declined to report the results. In this case, the data are informative and should be included in the analysis. However, there is no way to disambiguate case 1 from case 2, so all non-matched experiments had to be excluded. To evaluate how much this decision impacted on the results, we progressively injected the database with decrease experiments randomly sampled from the control dataset (see Data collection). Each one of these experiments was matched with a dummy increase experiment with no foci. We added 10 of these dummy pairs of experiments to the main database at each time, until we injected 220 dummy pairs, and repeated this procedure 30 times. The same analysis was made with increases. If the results hold to a consistent injection of dummy-pairs in the database, we could conclude that our network do not suffered the decision to exclude the unmatched experiments.

## Results

Our first selection of studies produced 85 decrease and 85 increase experiments coupled to each other. Table 1 contains the number of experiments for each of the disorders of such dataset. The distribution of psychiatric and neurological pathologies in those experiments was asymmetrical, with 64 experiments related to 9 psychiatric diagnoses and 21 experiments to 4 neurological ones. Of the latter, 13 were related to epilepsy, the remaining to extrapyramidal and movement disorders (*n* = 4), Parkinson’s disease (*n* = 2) and dystonia (*n* = 2). Thus, it seems that the coupling of decreases and increases is more characteristic of psychiatric conditions than neurological ones (with the only exception of the epilepsy), therefore, we decided to focus our analysis only on psychiatric disorders. The COA-O network comprising neurological disorders can be seen at Supplementary Fig. S2.

### Decrease-increase association network

The COA-O network for psychiatric diseases is presented in Figure 2. Although our network-mapping procedure produced 355 nodes, only 97 of them (32%) showed statistically significant co-occurrent changes with another region. For this subset of 97 nodes, we identified 292 significant co-occurrence probabilities, which represent edges in the COA-O network. Each edge in this network connects two areas showing GM changes in opposing directions. Of the 97 nodes in the network, 46 showed a GM decrease, 38 showed a GM increase, and 13 showed both (a node can assume both the roles of decrease and increase in different edges)..

**Figure 2:**
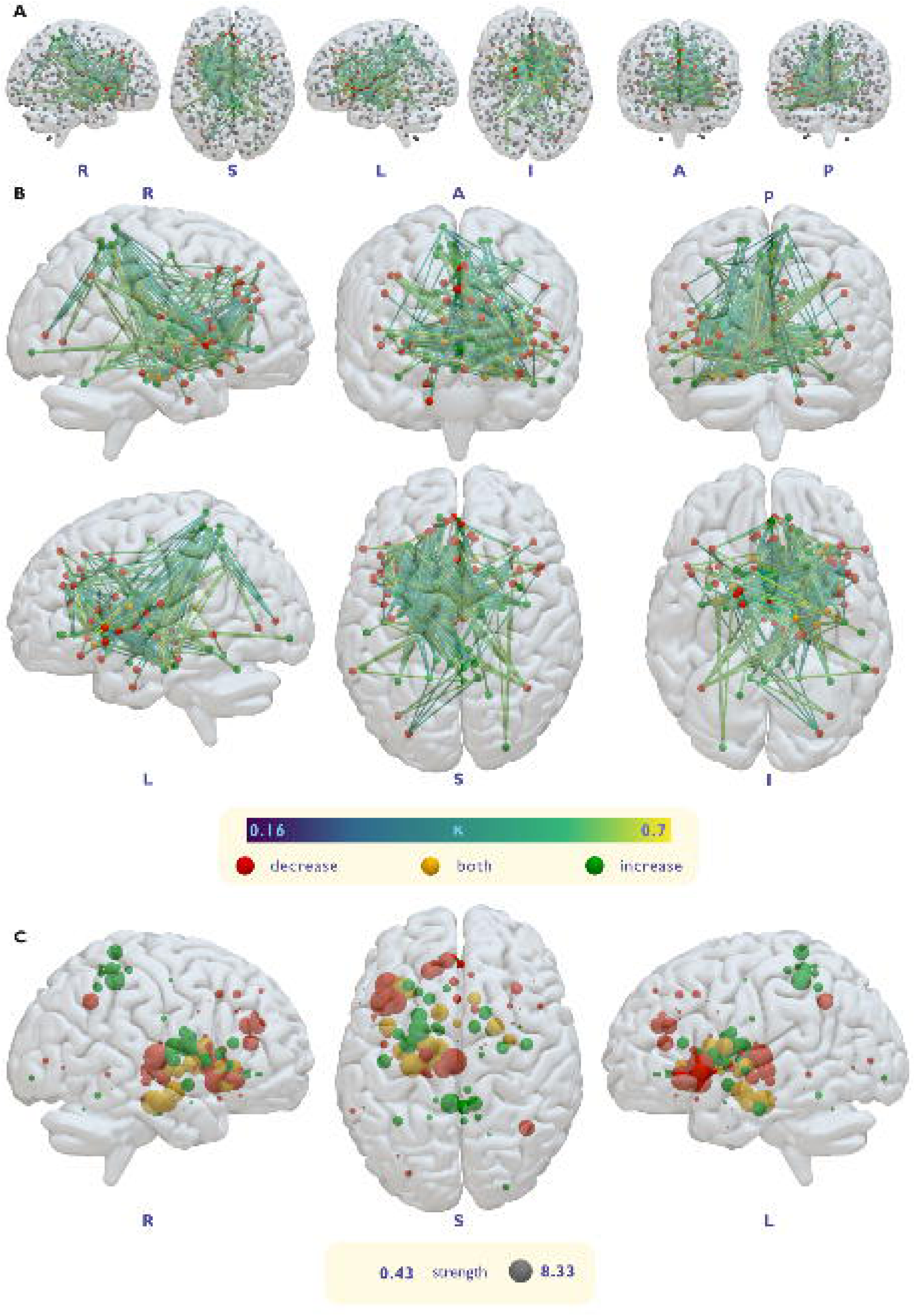
**A**: network of co-alterations of opposite GM changes, showing the unconnected nodes in grey color. **B**: network of co-alterations of opposite GM changes. **C**: nodal strength of the network. The size of the nodes is proportional to the weighted degree centrality.

GM increases and decreases are not uniformly distributed throughout the network. For instance, the superior parietal lobule (SPL) has a high concentration of areas with increased GM (0 nodes showing a decrease and 6 showing an increase), while the left insula and left inferior frontal gyrus (IFG) contain many nodes of decreased GM (10 decresase nodes, 1 increase node, 3 nodes are both decrease and increase), as showed by their nodal strength. The strengths (i.e. weighted degree centrality) of decreases and increases were calculated as the sum of each row and each column of the COA-O matrix, respectively. Node strength is unevenly distributed between hemispheres, with the left hemisphere having higher strength nodes than the right (Fig. 2 B). In fact, of the 25 nodes with highest strength, only 5 were in the right hemisphere (Table 4). Involvement of regions in the occipital lobe and posterior temporal lobes, and anterior prefrontal cortex, in the network is sparse. Most of the nodes, especially the ones with high strength, are situated in the anterior temporal and inferior frontal cortices, thalamus, and basal ganglia.

**Table 4:** the 25 nodes with the highest strength

| Node | x | y | z | Type of alteration | Strength |
| --- | --- | --- | --- | --- | --- |
| Left BA 13 | -42 | 16 | 0 | decrease | 8.33 |
| Left Thalamus | -2 | -18 | 6 | decrease | 7.74 |
| Left BA 47 | -40 | 26 | -6 | decrease | 7.35 |
| Left BA 35 (Parahippocampal gyrus) | -26 | -20 | -20 | both decrease and increase | 7.2 |
| Left BA 47 | -36 | 16 | -8 | decrease | 6.89 |
| Left Putamen | -22 | 0 | 12 | increase | 6.47 |
| Left BA 28 | -14 | -16 | -20 | both decrease and increase | 6.36 |
| Left Amygdala | -18 | -8 | -14 | both decrease and increase | 6.27 |
| Left BA 5 | -4 | -42 | 50 | increase | 5.94 |
| Left Thalamus | -8 | -22 | -2 | decrease | 5.63 |
| Left BA 36 (Parahippocampal gyrus) | -32 | -12 | -20 | both decrease and increase | 5.59 |
| Left Putamen | -28 | -8 | 12 | increase | 5.36 |
| Left Thalamus | -10 | -10 | 0 | both decrease and increase | 5.23 |
| Left BA 13 | -40 | 2 | 12 | both decrease and increase | 5.23 |
| Right Putamen | 22 | 2 | 2 | both decrease and increase | 5.18 |
| Left Caudate Body | -10 | 10 | 12 | both decrease and increase | 5.17 |
| Left BA 32 | -16 | 34 | 24 | decrease | 5.03 |
| Left Putamen | -22 | 8 | 6 | increase | 5.01 |
| Left BA 9 | -6 | 40 | 24 | decrease | 4.85 |
| Right BA 40 | 40 | -56 | 38 | decrease | 4.83 |
| Left BA 32 | -12 | 36 | 14 | decrease | 4.83 |
| Left BA 47 | -32 | 26 | 0 | both decrease and increase | 4.82 |
| Right BA 5 | 10 | -42 | 66 | increase | 4.57 |
| Right BA 5 | 4 | -44 | 56 | increase | 4.34 |
| Left BA 47 | -28 | 22 | -10 | both decrease and increase | 4.29 |

These findings were replicated using the Brainnetome atlas, producing a similar network and thus demonstrating that our method is independent of the node definition procedure (Fig. S4).

### Relationship between co-occurrent changes and functional connectivity

Contrary to our first hypothesis, the 292 significant Patel’s κ values of the COA-O matrix were not correlated with the 292 corresponding Pearson’s r values of the FC matrix (*r* = −0.07, non-significant at p=0.05). If COA-O edges were correlated to FC, it could be expected that most of them connected nodes belonging to a same canonical resting state networks (RSN), within which FC values tend to be high. Given the lack of correlation between our COA-O edges and normative FC, we hypothesised that the edges of the COA-O network tend to connect nodes belonging to different RSNs. Thus, each node was assigned to one of the RSN of the Yeo7 parcellation (Yeo *et al*., 2011), plus the cerebellum and the Thal/BG. Of the 292 edges, only 49 are between nodes of the same RSN, whereas 83% link nodes that belong to different RSNs. In comparison, COA-I and COA-D networks have a lower fraction of between-RSNs edges. The between-RSN edges of the COA-D network represent 78% of the total; the fraction is 68% for the COA-I. To evaluate the statistical significance of these differences, we randomized the COA-O, COA-D and COA-I networks using the Maslov-Sneppen algorithm (Maslov and Sneppen, 2002; Rubinov and Sporns, 2010) to preserve the degree distribution of each network. We then computed the differences in between-RSN edge fractions between the randomized networks and repeated the process 1000 times to build an empirical null distribution. The observed between-RSN fractions were significantly higher for the difference between the COA-O network and COA-D network (p<0.001) and between the COA-O network and COA-I network (p<0.001). Thus, although in comparison with with the COA-D network the difference is only of 5% (13% with the COA-I network), co-occurrent GM changes of opposing polarity are significantly more likely the occur between brain regions belonging to different functional networks.

(Maslov and Sneppen, 2002; Rubinov and Sporns, 2010)Focusing on the COA-O network, many between RSN edges are incident upon DMN nodes (Fig. 3). Most nodes mapped to the DMN show decreased GM, thus the areas to which these nodes connect in the COA-O network almost always show increased GM (especially Thal/BG, salience andexecutive control networks). The DMN is also the RSN with more nodes in the COA-D network, but it is poorly represented in the COA-I network, further supporting a higher frequency of GM reductions in this brain network. Almost all COA-O edges within the DMN connect to a single node showing increased GM located in the ventromedial prefrontal cortex ({x,y,z} = [-16, 34, -4]). This indicates that is is relatively rare to find co-occurring GM increases and decreases within the DMN.

**Figure 3:**
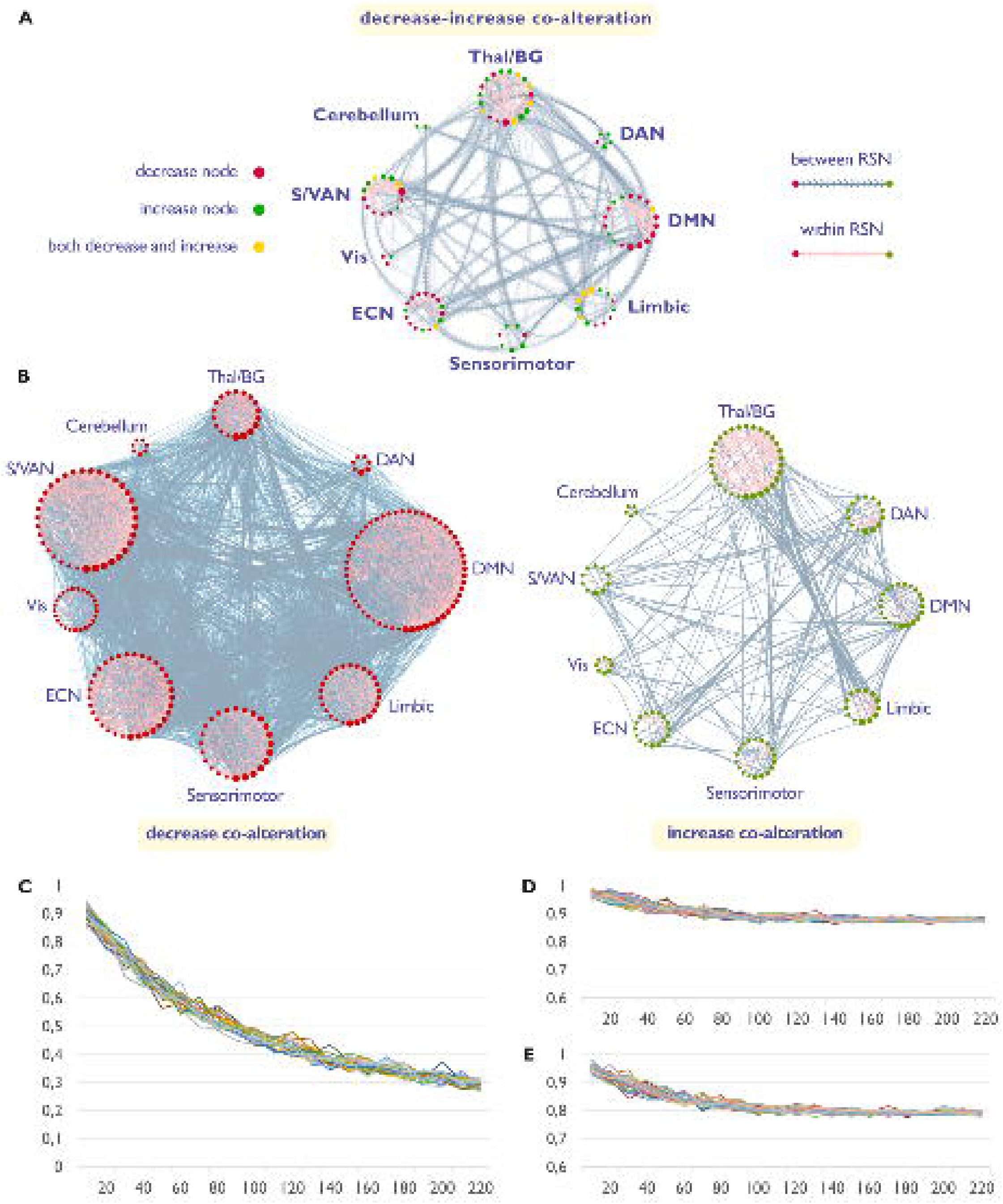
**A** and **B**: co-alteration networks represented in 2D, dividing each node for one of the Yeo7 RSN, plus cerebellum and thalamus/basal ganglia. The size of the nodes is proportional to the degree centrality, standardized across networks. DAN: dorsal attention network; DMN: default mode network; ECN: executive control network; Vis: visual network; S/VAN: salience/ventral attention network; Thal/BG: thalamus and basal ganglia, comprising upper midbrain. **A**: Decrease-increase co-alteration network. The transparency of the edge is inversely proportional to its κ value. Note that the arrows do not mean that the network is directed in a strict sense as the Patel’s κ is not a measure that provide a directionality. However, each edge links two nodes of different modality: one is a decrease and the other is an increase. Arrows were used to indicate at which side of the edge is the increase node. **B**: decrease only and increase only co-alteration networks. **C**: plot of the 30 runs of the Fail-safe analysis. On the x-axis: level of the null model. For each level, 10 random modelled alteration maps were added to the database. On the y-axis: values of Pearson’s r between the original decrease-increase association network and each of the level of the model. **D**: plot of the 30 runs of the dummy-pairs analysis with control decreases plus dummy increases. **E**: plot of the 30 runs of the dummy-pairs analysis with control increases plus dummy decreases.

As with the DMN, the ECN and, overall, the SN mostly comprise nodes showing GM decreases. In contrast, the dorsal attention network (DAN) is comprised almost only of nodes showing GM increases, located in the SPL (Fig. 2), and mostly associated with decreases in the Thal/BG and IFG (Fig. 2, Fig. 3 and Table S3). The cerebellum has only two increase nodes associated to the DMN and the ECN and that were not replicated using the Brainnetome atlas. In general, most edges of the COA-O are focused on regions in the telencephalon. A table listing all edges can be found in the Supplementary Materials (Table S3).

The node distribution across RSN is non-random. The DMN and Thal/BG are the RSNs with more nodes, while the cerebellum and visual network are the less represented. To evaluate if this spatial distribution is statistically significant, we randomly picked 97 nodes from a homogeneous parcellation (Fornito *et al*., 2010; Zalesky *et al*., 2010). A permutation test showed that, apart from the DMN, all the RSNs have a number of nodes significantly different from the null model (*p* = 0.05, one-tailed t-test, 10000 permutations). RSNs with few nodes have less nodes than expected from chance (Visual, *p* < 0.001; Sensorimotor, *p* = 0.0398 ; DAN, *p* = 0.0119; Cerebellum, *p* < 0.001), while the bigger ones have significantly more nodes (Salience, *p* = 0.0105; Limbic, *p* = 0.0326, Executive control, *p* = 0.0381; Thal/bg, *p* < 0.001).

### Fail-safe and dummy-pairs analyses

As our COA-O dataset of experiments is relatively small, we assessed the robustness of our results to the injection of noise with a modified Fail-safe technique (Acar *et al*., 2018). We observed that the COA-O network is still correlated with *r* ≈ 0.3 even after we added 220 null maps (∼350% of the original dataset, Fig. 3 C). This means that, if our dataset was much larger, the network might remain relatively stable, unless the injection of real data was somehow much more harmful to the results than that of null studies.

To verify that our results were not severely affected by the necessity of including only the couple of experiments showing opposing GM changes, we added a set of uncoupled experiments, paired with an empty one. The results can be seen at Fig. 3 D and E. The network resulting from the addition of 220 decrease experiments paired with empty ones correlates with the original one at r=0.88 (averaged across 30 runs). Applying the same procedure adding only GM increase experiments produced a network that correlated with the original one, on average, r=0.79. Thus, our results would if we did not exclude non-matching experiments.

### Directionality of the decrease-increase associations

We further assessed the directionality of the edges of the COA-O network using the Patel’s τ, which compares the conditional probabilities of having an alteration of region A given a change in B and of a change in B given a change in A. A positive edge means that it is more probable to have an increase given a decrease than the other way round, while a negative edge means that it is more probable to have an increase given a decrease. The τ network is characterized by both positive and negative edges (Fig. 4) although positive edges are slightly stronger and more numerous. Many negative edges are incident upon nodes of the parietal lobe that are characterized by GM increases. Conversely, the nodes in the left IFG and the thalamus, which represent areas of GM decrease, are connected by many positive edges (Fig 4 and S5). Many positive edges link a decrease of the salience network or the DMN to a decrease in the Thal/BG, but also DMN to SN and SN to limbic network. Negative edges are more distributed across RSNs (Fig. 4). In general. these findings indicate that the edges of the COA-O network are often directed from the decrease nodes to the increase nodes, but also the opposite can be true.

**Figure 4:**
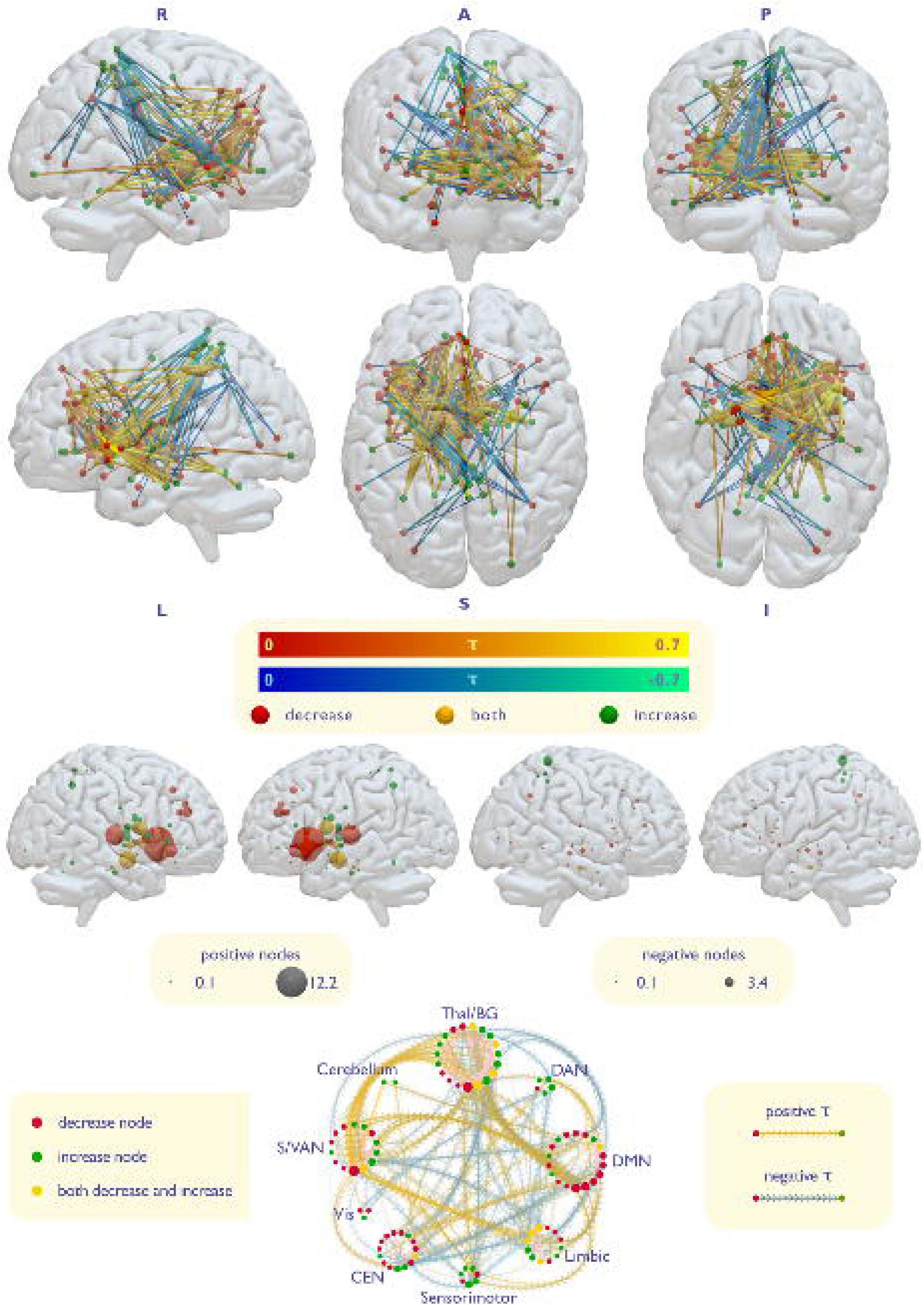
**A**: directed network of decrease-increase co-alteration. Positive edges indicates that the decrease node is dominant on, and possibly influences, the increase node. Negative edges indicates the opposite. **B**: hubs of the directed network, divided for positive hubs (left) and negative hubs (right). Note that these images were produced calculating the strength of each node on the whole Patel’s τ matrix, and then separated between the nodes with a positive value and those with a negative value. Thus, the strength of each node represents its balance between the positive and negative edges incident upon it. Therefore the size of the node is not proportional to the number of its edges, since positive and negative edges incident upon a same node cancel each other. See Supplementary Figure S5 for a representation of the strength of positive-only and and negative-only edges. **C**: 2D representation of the directed network. Positive edges indicates that the decrease node is dominant on, and possibly influences, the increase node. Negative edges indicates the opposite. DAN: dorsal attention network; DMN: default mode network; ECN: executive control network; Vis: visual network; S/VAN: salience/ventral attention network; Thal/BG: thalamus and basal ganglia, comprising upper midbrain.

## Discussion

Our meta-analytic network mapping procedure evaluated the statistical relationship between GM increases and decreases across neurological and psychiatric disorders. Co-occurrent decreases and increases in GM volume are rarely reported in neurological diseases, with epilepsy being an exception. Conversely, such co-occurrent changes where more common in psychiatric disorders. This result might reflect a reporting bias in the neurology literature to focus only on GM decreases,However, it may also represent a genuine increase in the likelihood of observing GM increases in psychiatric disorders and epilepsy. In fact, a common thread linking these two is that many of them have a neurodevelopmental origin, which may provide greater opportunity for plastic adaptations and thus the emergence of GM increases.

In psychiatric disorders, our analysis revealed that GM increases and decreases are not independent; instead, many areas show a GM change that is statistically related to a change of opposite polarity in other areas. The resulting COA-O network presents a series of interesting features. For instance, there appears to be a preponderance of left hemisphere nodes being more strongly involved in such coordinated GM changes (Fig. 2). The left hemisphere dominance in the COA-O network is intriguing and might suggest a differential involvement of the two hemispheres in the anatomy of psychiatric disorders. This observation is reminiscent of the recent finding that the hubs of the COA-D are symmetric across the hemispheres to those of the COA-I network (Cauda *et al*., 2019*a*), in the fact that they both highlight how the two hemispheres might be differently involved by opposing GM alterations.

The nodes involved in the COA-O phenomenon are not randomly distributed across the networks. In particular, the SN and ECN showed more nodes than expected by chance, consistent with their presumed importance in psychiatric pathology (Palaniyappan and Liddle, 2012; Goodkind *et al*., 2015; McTeague *et al*., 2016, 2017; Sheffield *et al*., 2017; Sha *et al*., 2019). The thalamus and basal ganglia also showed more significant nodes than expected by whereas the DMN does not. In fact, most canonical RSNs contained significantly fewer COA-O nodes than expected by chance. Notably, while the Thal/BG showed an approximately equal number of decrease and increase nodes, the DMN is mostly characterized by decreases. Such decreases were often found to be associated to Thal/BG increases, and increases in the the SN and ECN. The ECN and, to a lesser extent, the SN, show a similar propensity for decrease nodes associated with Thal/BG increases, indicating that GM decreases in the three higher order cortical networks are often accompanied by GM increases in subcortical structures.

Another relevant aspect of the COA-O network was the tendency of its edges to connect different RSNs. In fact, analysing the COA-O network for psychiatric disorders, we observed that it is not correlated to FC, and that its edges tend to connect different RSNs. Prior work has shown a closer association between FC and co-occurrent GM decreases or increases, which can be largely explained by models of diffusion along connectivity pathways (Cauda *et al*., 2018*b*; Raj and Powell, 2018). Our findings thus suggest that co-occurent GM changes of opposing polarity may emerge through a distinct phenomenon. Candidate mechanisms include effects of medication, direct effects of disease, or compensatory or maladaptive responses to insult.

### GM increases due to medication

It is possible that increased GM in psychiatric disorders is not due to the disease process, but instead reflects a secondary consequence of medication. For instance, it has been reported that lithium, commonly used to treat bipolar disorder, has a neurotrophic and morphometric increase effect (Manji *et al*., 1997, 1999; Moore *et al*., 2000; Chen *et al*., 2000; Fukumoto *et al*., 2001; Sassi *et al*., 2002; Angelucci *et al*., 2003; Hashimoto *et al*., 2003; Beyer *et al*., 2004; Frey *et al*., 2006; Bearden *et al*., 2007; Monkul *et al*., 2007; Yucel *et al*., 2007; Bearden *et al*., 2008; Yucel *et al*., 2008; Kempton *et al*., 2008; Hammonds and Shim, 2009; Lyoo *et al*., 2010; Hajek *et al*., 2012; López-Jaramillo *et al*., 2017; Hibar *et al*., 2018). Similarly, the use of conventional antipsychotics has repeatedly been associated with basal ganglia (Muller and Seeman, 1977; Chakos *et al*., 1994, 1998, Keshavan *et al*., 1994*b*; Murali *et al*., 1995; Sedvall *et al*., 1995; Shihabuddin *et al*., 1998; Kippin *et al*., 2005; Chopra *et al*., 2020)(Kippin *et al*., 2005) and thalamic (Gur *et al*., 1998; Strungas *et al*., 2003; Dazzan *et al*., 2005) anatomic increases.

Given that many patients included in our meta-analysis were undergoing medication at the time of the scan (Table S4), this variable is likely to have some impact on the COA-O network. However, there is some evidence that suggests that medication alone is unable to explain our results: first, morphological effects of medication are often found to be localized to restricted regions, such as medial temporal lobe and subgenual cortex with lithium (Germaná *et al*., 2010; Hafeman *et al*., 2012) and BG with antypsichotics (Navari and Dazzan, 2009), while our increase nodes are situated in many other brain areas; second, some studies report that medications could attenuate pathological decreases rather than increase GM volume in patients compared to controls (Sheline *et al*., 2003; Wada *et al*., 2005; Hibar *et al*., 2016; Zung *et al*., 2016; Sarrazin *et al*., 2019); third, atypical antipsychotics, although found by some to be neurotrophic and induce neurogenesis (Wakade *et al*., 2002; Bai *et al*., 2003; Halim *et al*., 2004; Wang *et al*., 2004; Park *et al*., 2006) produced mixed volumetric findings (Massana *et al*., 2005; Navari and Dazzan, 2009) but often no increase effects were found in the BG (Chakos *et al*., 1995; Frazier *et al*., 1996; Westmoreland Corson *et al*., 1999; Lang *et al*., 2001, 2004; Scheepers *et al*., 2001). Moreover, a study reports an absence of increased volume of BG also for typical antipsychotics (Kreczmanski *et al*., 2007); fourth, anticonvulsant drugs, used in the treatment of bipolar disorder but also for epilepsy (the only neurologic disease with a non-negligible number of experiments in our database) showed to produce decreases or no effect (Chang *et al*., 2009; Germaná *et al*., 2010; Abé *et al*., 2016; Hibar *et al*., 2018); fifth, the increased striatal volume of relatives of schizophrenic patients suggests a genetic factor (Oertel-Knöchel *et al*., 2012). Similarly, increased cortical thickness and subcortical volume were found also in drug-naive patients with depression (Qiu *et al*., 2014; Reynolds *et al*., 2014; Yang *et al*., 2015; Zuo *et al*., 2018; Li *et al*., 2019; Suh *et al*., 2019); sixth, patients with autism or development disorders included in our study were not under drug treatment (Table S4). Therefore, it seems that medications alone are unable to explain the phenomenon of DIA; seventh, if GM increases were purely explained by medication, they would most likely be statistically associated to all, or most of, the GM decreases. On the contrary, the nodes of increase were selectively co-altered with one or few decreases. The effect of medication thus cannot provide the sole explanation for the observed findings in the COA-O network.

### Co-occurrent increases and decreases as a direct effect of the pathology

One possible hypothesis might be that GM increases are as direct effect of the disease process. Co-occurring GM decreases and increases could arise due to regionally distinct effects of inflammatory processes (Li *et al*., 2019), or, possibly, altered developmental pruning (Keshavan *et al*., 1994*a*). Another option could be that, due to dysregulation of ascending neuromodulatory projection systems (Davis *et al*., 1991; Sesack and Carr, 2002), different areas of the brain may become hyperactivated or hypoactivated, potentially resulting in coordinated and concomitant increases and decreases in volume through activity-dependent plasticity. These changes could be driven by neurodevelopmental miswiring of connectivity, resulting in a de-differentiation of function (Fornito *et al*., 2015).

### Co-occurrent increases and decreases as a form of compensation

Another hypothesis might be that GM increases reflect a compensatory response to decreases. In fact, it has been reported that the brain topology of chronic patients shows modifications that appears to counter or normalize those occurred as consequence of a pathologic perturbation (Lord *et al*., 2012; Hillary *et al*., 2015). The seemingly most logical consequence of this view should be that the COA-O would happen between regions with similar function and thus belonging to the same RSN. Indeed, it was previously shown that, in schizophrenic patients, over- and under-activations are often located in topologically close areas (Crossley *et al*., 2016). Our data are not consistent with this scenario, as COA-O edges are not correlated with FC and are more often between-than within-RSN (Fig. 2). Given prior reports that strongly connected areas tend to share GM changes in the same direction (Cauda *et al*., 2018*b*; Shafiei *et al*., 2019), our findings suggest that such regions may have a limited capacity for compensation, possibly due to diaschisis and other maladaptive responses (Carrera and Tononi, 2014; Fornito *et al*., 2015). Therefore, a region which is not strongly connected to a damaged area but has a related function may be the best candidate to take on a compensatory role. For instance, the SPL nodes of increases are especially associated with decreases in the insula/IFG and Thal/BG. Despite belonging to distinct RSNs, these areas have all been implicated in pain perception (Chudler and Dong, 1995; Strigo *et al*., 2003; Fitzek *et al*., 2004; Lu *et al*., 2004; Kong *et al*., 2006; Schoedel *et al*., 2008; Freund *et al*., 2009; Tseng *et al*., 2010; Bräscher *et al*., 2016), somatosensory perception in general (Olausson *et al*., 2002, 2005; Yoo *et al*., 2003; Stoeckel *et al*., 2004; Nagy *et al*., 2006; Robbe, 2018), and motor functions (Stephan *et al*., 1995; Kertzman *et al*., 1997; Binkofski *et al*., 1999; Jovicich *et al*., 2001; Herrero *et al*., 2002; Groenewegen, 2003; Lotze *et al*., 2006; Beurze *et al*., 2007; Naito *et al*., 2008; Langner *et al*., 2014), although the SPL can also be associated to top-down attention as part of the DAN (Corbetta and Shulman, 2002; Yeo *et al*., 2011).

The concept of compensation usually involves an adaptive change for the individual, however some have suggested that neuroplastic modifications consequential to psychiatric diseases might also be dysfunctional for the patient well-being (Amad *et al*., 2019; Palaniyappan, 2019).

### Directionality of the associations

Using Patel’s τ, we also produced a directed network of COA-O, representing the ascendancy (influence) of decreases to increases. The resulting network shows a clear separation of positive and negative edges, with the former being more prevalent between fronto-temporal nodes and the latter being incident upon parietal nodes (Fig. 4). Although this is an interesting observation that suggests that the COA-O mechanisms might work differently in distinct parts of the brain, there are several possible explanations for this observation. Critically, while the unbalance of conditional probabilities might indicate an influence of a node on the other, it could result also from the action of a third agent. In fact, if node *a* and *b* are both influenced by an element external to the couple, but one of the two nodes is more vulnerable to its action than the other, we would obtain the same unbalanced probabilities of alteration. Such a third agent could be another node or the pathology itself, to which the two regions would be differently vulnerable. Such differential vulnerability expresses in two ways: one of the nodes undergo a GM decrease while the other an increase, and one of the two node is more likely to be modified than the other. Therefore, the ascendancy as we calculated it could be either interpreted as a measure of influence of a node on the other, or as an evidence of a sort of primacy of a node over the other in the system of COA-O. In fact, considering that the COA-O edges do not correspond to those of FC, the hypothesis of a direct influence of a node on another seems the less likely of the two.

### Limitations and future directions

The main limitation of this study comes from a relatively small sample of included experiments. Although we adopted a transdiagnostic approach, only 64 comparisons reported both increases and decreases on psychiatric patients in the BrainMap database. We suspect increases can be sometimes overlooked and not reported by some authors, producing a file-drawer effect. However, our Fail-safe technique showed that our network remains relatively preserved also after the injection of ∼350% of noise, reassuring us about the validity of our results. Moreover, even if we include non-coupled experiments, the network remains remarkably similar, indicating that the COA-O edges can be attenuated, but not radically changed if including papers that do not presented both GM changes in the database.

The transdiagnostic approach embraced here was motivated by the interest in general brain mechanisms of COA-O, but disease-specific investigations would be of great interest as well. The scarcity of experiments retrieved by our search prevented us to do so, but future meta-analyses might be able to gather more data.

Longitudinal or cross-sectional studies are also needed to ascertain the presence of COA-O in individual’s brains Moreover, functional neuroimaging and clinical assessment would be critical to evaluate the eventual compensatory function of the COA-O phenomenon.

The use of the Patel’s τ has been advocated by some as a valid method to investigate network directionality (Smith *et al*., 2011), but its use in effective connectivity has also been severely questioned by others (Wang *et al*., 2017). However, our study did not used the Patel’s τ in search of a strict causal directionality, but to investigate a more general notion of directionality between nodes.

## Conclusions

Our analysis provides evidence for coordinated GM decreases and increases in psychiatric disorders, with such coordinated changes principally affecting higher-order association networks and subcortical regions. Our findings (Adachi *et al*., 2012)open a new line of research into the mechanisms underlying coordinated GM changes of opposing polarity, and we hope that they draw greater attention on the often-neglected GM increases observed in patients.

## Competing interested

The authors declare no competing interest.

