## Supplementary Materials for "A meta-analytic approach to mapping co-occurrent grey matter volume increases and decreases in psychiatric disorders"

Fig. S1: Prisma Flow chart of the selection of studies.

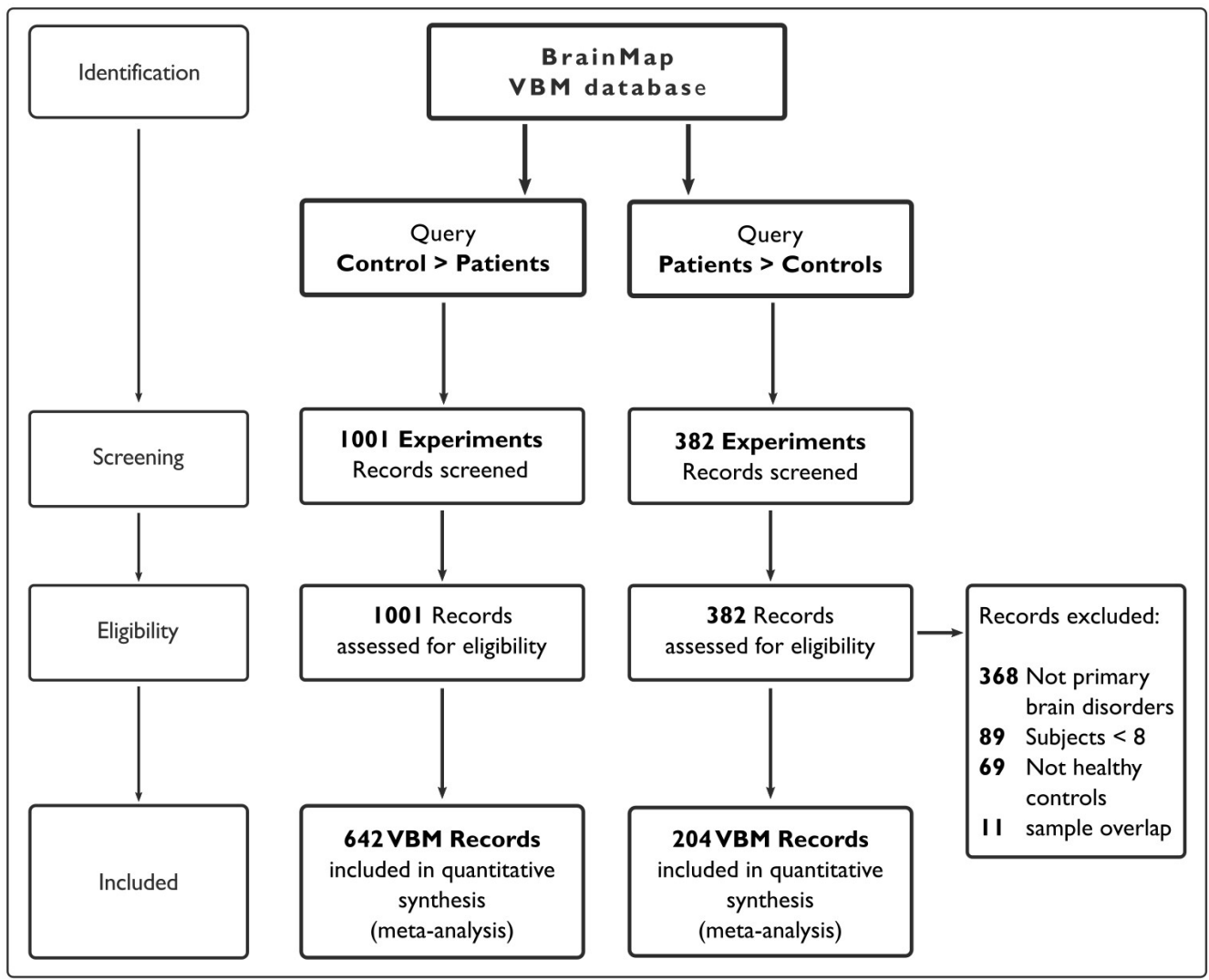

**Fig S2:** network of co-alterations of opposite GM changes calculated with both psychiatric and neurological diseases.

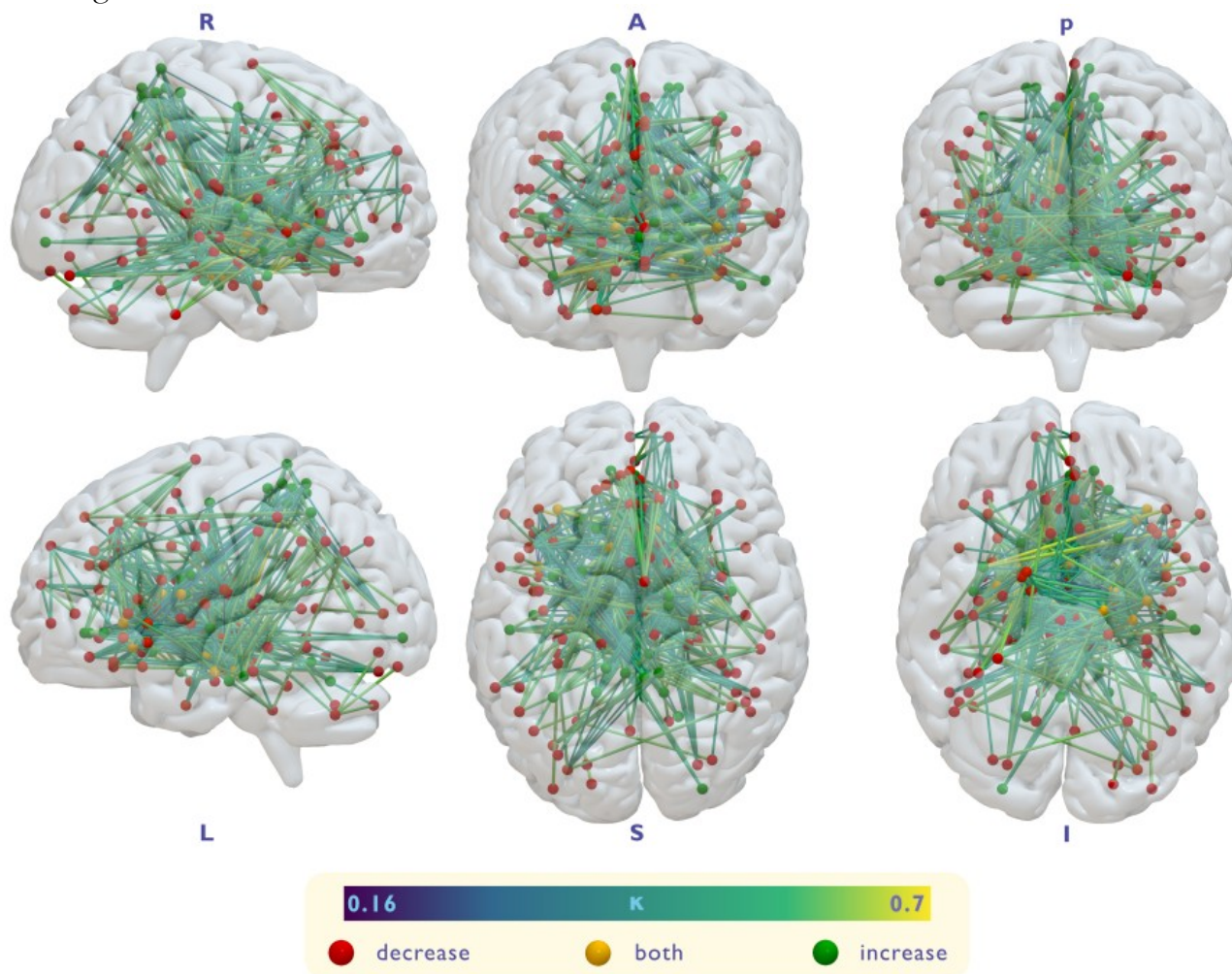

**Fig. S3:** matrix of co-alteration of GM decreases and increases.

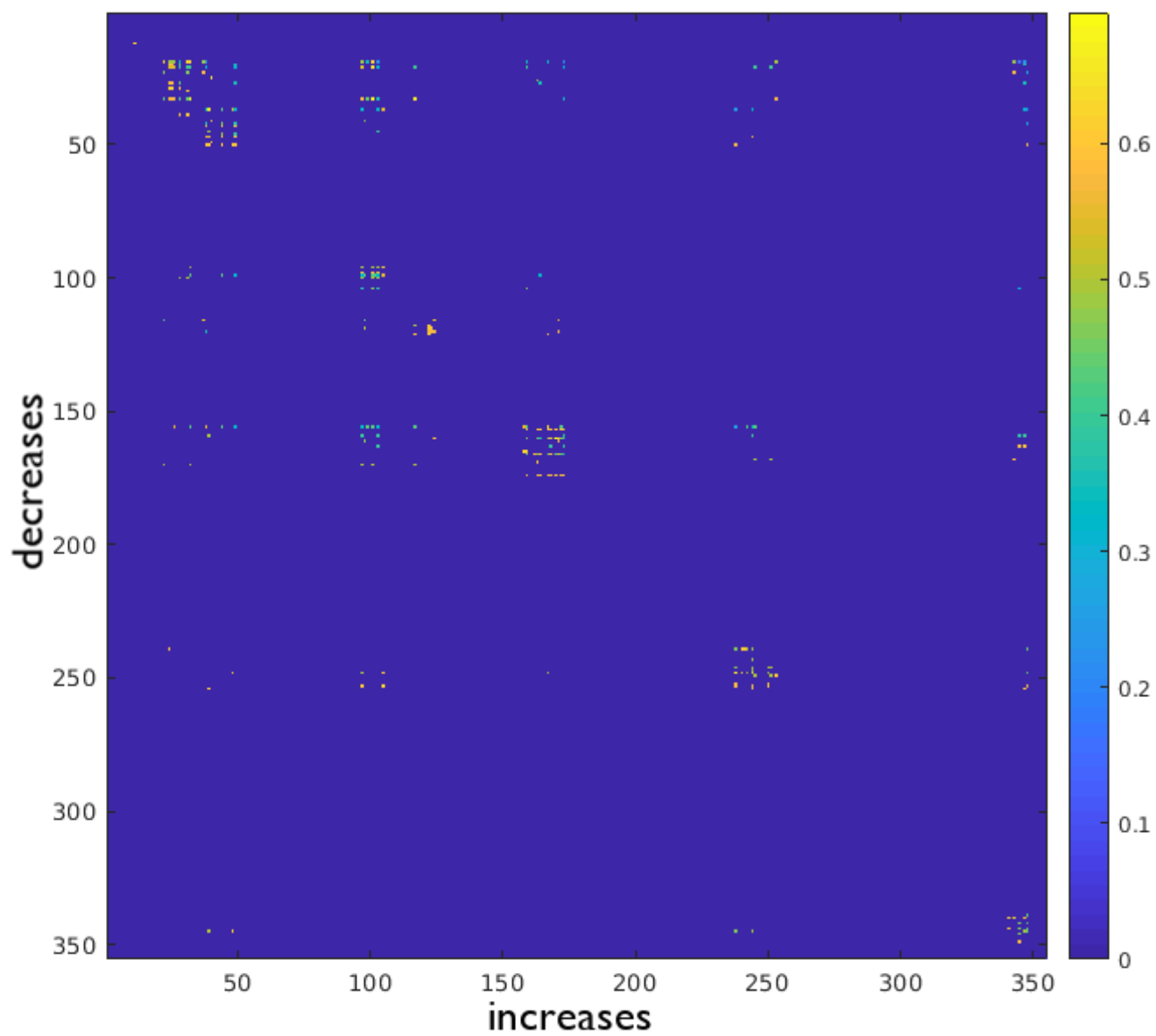

**Fig S4:** network of co-alteration of opposing GM changes calculated with the Brainnetome Atlas.

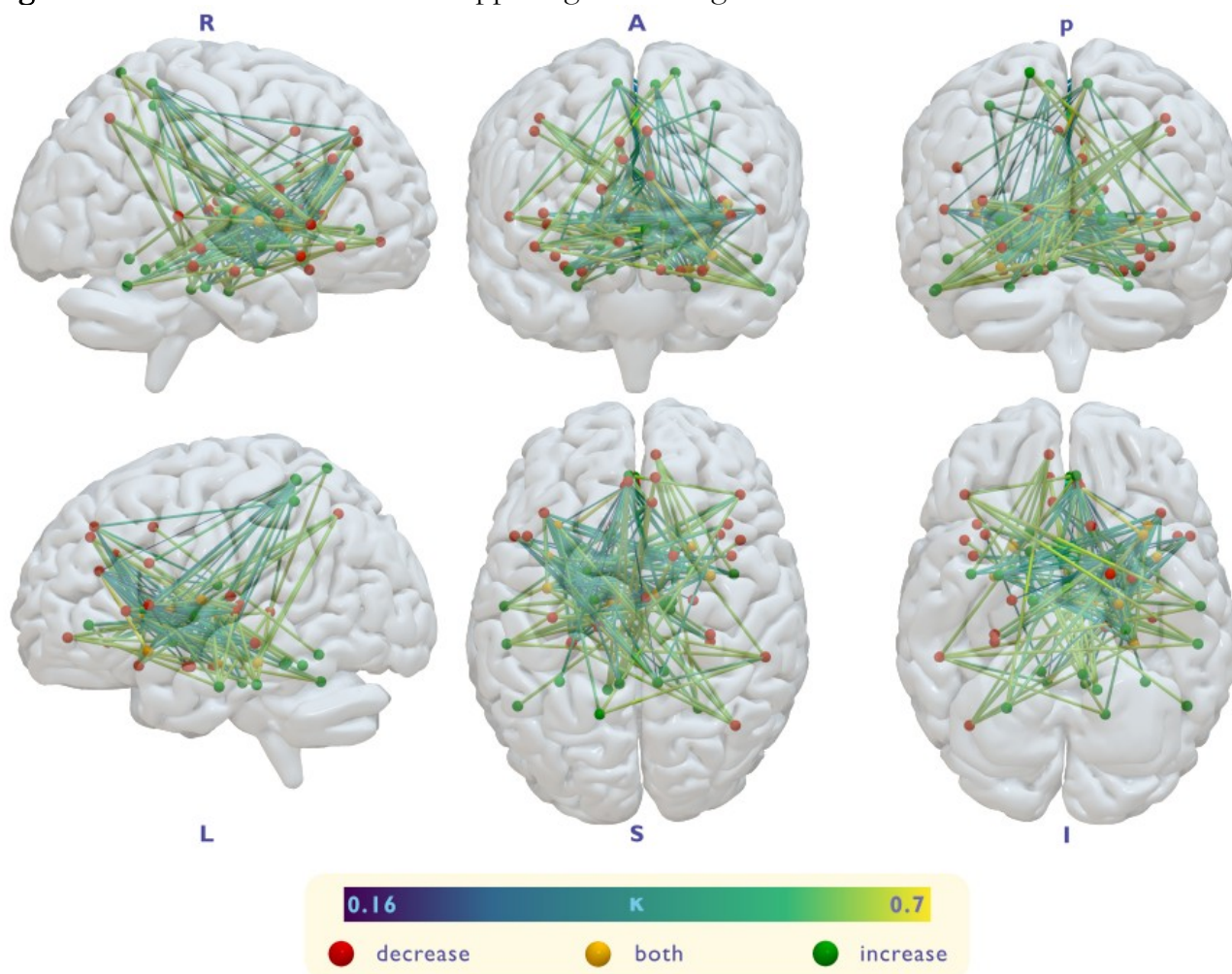

**Fig S5:** strenghts of the nodes of the directed network calculated only on the positive nodes (left) and negative nodes (right).

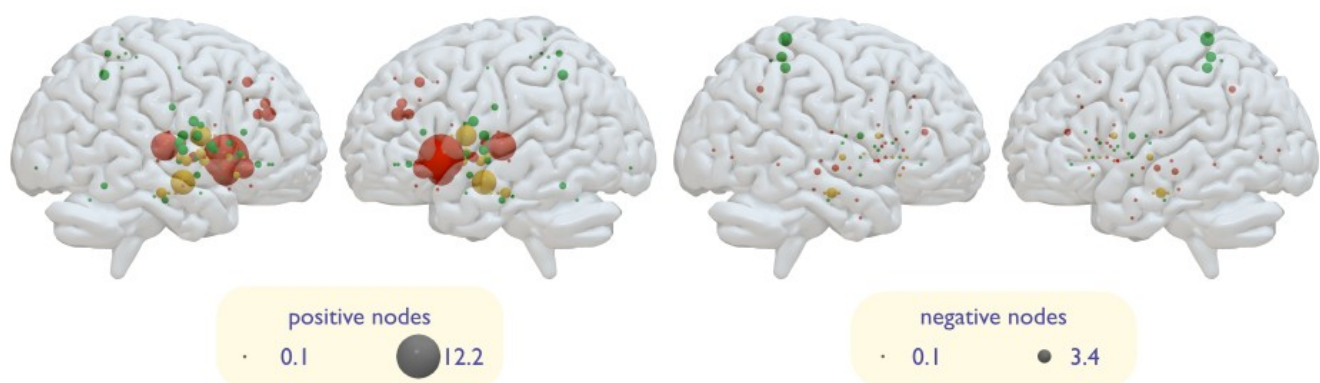

**Table S1:** experiments included in the control meta-analysis of decrease-only co-alteration.

| AUTHOR | CONDITION | # SUBJECTS | ICD-10 CODE | DIAGNOSIS |
| --- | --- | --- | --- | --- |
| AnanthH,2002 | Normals>SchizophreniaPatients,GrayMatterVolume | 20 | F20 | Schizophrenia SZ |
| AntonovaE,2005 | Normals>SchizophreniaPatients,GreyMatterVolume | 40 | F20 | Schizophrenia SZ |
| BassittDP,2007 | Normals>Schizophrenia,GrayMatterVolume | 30 | F20 | Schizophrenia SZ |
| BergÈD,2011 | Patients<Controls,Graymattervolume | 20 | F20 | Schizophrenia SZ |
| BonilhaL,2008 | SchizophreniaPatients<HealthyControls,GrayMatter | 13 | F20 | Schizophrenia SZ |
| BorgwardtSJ,2010 | MZtwinsconcordantforschizophrenia<controls,Grey | 28 | F20 | Schizophrenia SZ |
| BorgwardtSJ,2010 | MZtwinsdiscordantforschizophreniapatientsvs | 9 | F20 | Schizophrenia SZ |
| BoseSK,2009 | Patients<Controls,Greymattervolume | 33 | F20 | Schizophrenia SZ |
| CascellaN,2010 | HealthyControls>AllSchizophreniaPatients,Gray | 19 | F20 | Schizophrenia SZ |
| CascellaN,2010 | HealthyControls>DeficitSchizophreniaPatients,Gray | 19 | F20 | Schizophrenia SZ |
| CascellaN,2010 | HealthyControls>Non-deficitSchizophreniaPatients, | 31 | F20 | Schizophrenia SZ |
| Castro-ManglanoPD,2011 | Greymattervolumereduction,FEPpatients< | 20 | F20 | Schizophrenia SZ |
| ChowEW,2011 | 22q11.2DeletionSyndromePatients(withSchizophrenia< | 29 | F20 | Schizophrenia SZ |
| ChowEW,2011 | 22q11.2DeletionSyndrome(withSchizophrenia< | 29 | F20 | Schizophrenia SZ |
| ChuaSE,2007 | FEPPatients<Controls,Greymattervolume | 26 | F20 | Schizophrenia SZ |
| CookeMA,2008 | SchizophreniaPatients<HealthyControls,GrayMatter | 30 | F20 | Schizophrenia SZ |
| CuiL,2011 | Paranoid-typeSchizophreniaPatients<HealthyControls,Gray | 23 | F20 | Schizophrenia SZ |
| DengMY,2009 | Healthycontrolssecondassessment>Patients[re-scan | 10 | F20 | Schizophrenia SZ |
| DengMY,2009 | Healthycontrols>Patients[re-scanbeyond3weeks], | 10 | F20 | Schizophrenia SZ |
| DouaudG,2007 | AOS<Healthycontrols,greymattervolume | 25 | F20 | Schizophrenia SZ |
| EbdrupBH,2010 | SchizophreniaPatients(all)<HealthyControls,Grey | 38 | F20 | Schizophrenia SZ |
| EbdrupBH,2010 | SchizophreniaPatients(abs)<HealthyControls,Grey | 38 | F20 | Schizophrenia SZ |
| EbdrupBH,2010 | SchizophreniaPatients(non-abs)<HealthyControls,Grey | 38 | F20 | Schizophrenia SZ |
| EulerM,2009 | HealthyControls>SchizophreniaPatients,GrayMatter | 19 | F20 | Schizophrenia SZ |
| EulerM,2009 | HealthyControls>SchizophreniaPatients(Controllingfor | 19 | F20 | Schizophrenia SZ |
| FoongJ,2001 | ScizophreniaPatients<HealthyControls,Magnetization | 25 | F20 | Schizophrenia SZ |
| FoongJ,2001 | SchizophreniaPatients<HealthyControls,SignalIntensity | 25 | F20 | Schizophrenia SZ |
| Garcia-MartiG,2008 | Scz-AHPatients<HealthyControls,GreyMatter | 17 | F20 | Schizophrenia SZ |
| GiulianiNR,2005 | SchizophreniaPatients<HealthyControls,GrayMatter | 34 | F20 | Schizophrenia SZ |
| HaTH,2004 | ParanoidSchizophreniaPatients<HealthyControls,Gray | 35 | F20 | Schizophrenia SZ |
| HeroldR,2009 | HealthyControls>SchizophreniaPatients,GrayMatter | 18 | F20 | Schizophrenia SZ |
| HiraoK,2008 | HealthyControls>SchizophreniaPatients,GrayMatter | 20 | F20 | Schizophrenia SZ |
| HoneaRA,2008 | HealthyControls>SchizophreniaSpectrumDisorder,Gray | 169 | F20 | Schizophrenia SZ |
| HornH,2009 | SchizophreniaPatients<HealthyControls,GrayMatterVolume | 13 | F20 | Schizophrenia SZ |
| HornH,2010 | SchizophrenicPatients<HealthyControls,GrayMatterVolume | 20 | F20 | Schizophrenia SZ |
| HulshoffPolHE,2001 | Normals>SchizophreniaPatients,GrayMatter | 158 | F20 | Schizophrenia SZ |
| HulshoffPolHE,2004 | Normals>SchizophreniaPatients,GrayMatter | 158 | F20 | Schizophrenia SZ |
| HulshoffPolHE,2006 | HealthyControls>DiscordantforSchizophrenia, | 22 | F20 | Schizophrenia SZ |
| JanssenJ,2008 | SchizophreniaPatients<HealthyControls,GrayMatter | 25 | F20 | Schizophrenia SZ |
| JayakumarPN,2005 | FirstEpisodeSchizophreniavsHealthyControls,Gray | 18 | F20 | Schizophrenia SZ |
| JayakumarPN,2005 | FirstEpisodeSchizophreniavsHealthyControls,Gray | 18 | F20 | Schizophrenia SZ |
| KasparekT,2007 | Normals>SchizophreniaPatients,GrayMatterVolume | 18 | F20 | Schizophrenia SZ |
| KasparekT,2009 | Schizophreniapatients(GoodandPoorfunctioning)< | 18 | F20 | Schizophrenia SZ |
| KasparekT,2009 | PoorFunctioningFES<HealthyControls,Graymatter | 18 | F20 | Schizophrenia SZ |
| KasparekT,2009 | GoodFunctioning<HealthyControls,Graymattervolume | 18 | F20 | Schizophrenia SZ |
| KasparekT,2010 | First-EpisodeSchizophrenia<HealthyControls,Gray | 49 | F20 | Schizophrenia SZ |
| KawadaR,2009 | SchizophreniaPatients<HealthyControls,GrayMatter | 26 | F20 | Schizophrenia SZ |
| KawasakiY,2004 | SchizophreniaPatients<HealthyControls,GrayMatter | 25 | F20 | Schizophrenia SZ |
| KawasakiY,2004 | SchizotypalDisorderPatients<HealthyControls,Gray | 25 | F20 | Schizophrenia SZ |
| KawasakiY,2007 | Normals>SchizophreniaPatients,GrayMatterVolume | 30 | F20 | Schizophrenia SZ |
| KoutsoulerisN,2008 | HealthyControls>Schizophrenia,GreyMatter | 55 | F20 | Schizophrenia SZ |
| KoutsoulerisN,2008 | HealthyControls>SchizophreniaNegative,Grey | 59 | F20 | Schizophrenia SZ |
| KoutsoulerisN,2008 | HealthyControls>SchizophreniaPositive,Grey | 61 | F20 | Schizophrenia SZ |
| KoutsoulerisN,2008 | HealthyControls>SchizophreniaDisorganized,Grey | 55 | F20 | Schizophrenia SZ |
| KubickiM,2002 | Normals>SchizophreniaPatients,GrayMatterDensity | 16 | F20 | Schizophrenia SZ |
| LuiS,2009 | FamilialSchizophreniaProbands<HealthyControlProbands, | 10 | F20 | Schizophrenia SZ |
| LuiS,2009 | SporadicSchizophreniaProbands<HealthyControlProbands, | 10 | F20 | Schizophrenia SZ |
| LuiS,2009 | HealthyControls>First-EpisodeSchizophreniaPatients,Gray | 68 | F20 | Schizophrenia SZ |
| Marti-BonmatiL,2007 | SchizophreniaPatients<HealthyControls,Gray | 10 | F20 | Schizophrenia SZ |
| McIntoshAM,2004 | Normals>SchizophreniaPatients(SCZ),GrayMatter | 26 | F20 | Schizophrenia SZ |
| MedaSA,2008 | Normals>SchizophreniaPatients(JHU,MPRC,IOP,WPIC), | 9 | F20 | Schizophrenia SZ |
| MedaSA,2008 | HealthyControls>SchizophreniaPatients(JHU),Gray | 133 | F20 | Schizophrenia SZ |
| MedaSA,2008 | HealthyControls>SchizophreniaPatients(MPRC),Gray | 34 | F20 | Schizophrenia SZ |
| MedaSA,2008 | HealthyControls>SchizophreniaPatients(WPIC),Gray | 21 | F20 | Schizophrenia SZ |
| MeisenzahlEM,2008 | Normals>First-EpisodeSchizophrenia,GrayMatter | 93 | F20 | Schizophrenia SZ |
| MeisenzahlEM,2008 | Normals>RecurrentlyIllSchizophrenia,GrayMatter | 72 | F20 | Schizophrenia SZ |
| MeisenzahlEM,2008 | Normals>First-EpisodeSchizophrenia+Recurrently | 72 | F20 | Schizophrenia SZ |
| MeisenzahlEM,2008 | Normals>FESPatients,maskedbyNormals>REZ | 72 | F20 | Schizophrenia SZ |
| MeisenzahlEM,2008 | Normals>REZpatients,maskedbyNormals>FES | 72 | F20 | Schizophrenia SZ |
| MolinaV,2010 | FirstEpisodeSchizophreniavs.HealthyControls,Gray | 30 | F20 | Schizophrenia SZ |
| MolinaV,2011 | SZ<CS,GrayMatterVolume | 24 | F20 | Schizophrenia SZ |
| MolinaV,2011 | Healthycontrols>Patients,reductionsingreymattervolume | 30 | F20 | Schizophrenia SZ |
| MoorheadTW,2005 | HealthyControls>SchizophreniaPatients,GrayMatter | 25 | F20 | Schizophrenia SZ |
| NeckelmannG,2006 | SchizophreniaPatients>HealthyControls,GreyMatter | 12 | F20 | Schizophrenia SZ |
| NeckelmannG,2006 | CorrelationbetweenBPRSHallucinationScoresGrey | 12 | F20 | Schizophrenia SZ |
| O'DalyO,2007 | SchizophreniaPatients<HealthyControls,Greymatter | 28 | F20 | Schizophrenia SZ |
| OhnishiT,2006 | Control>Schizophrenia,GreyMatterVolume | 19 | F20 | Schizophrenia SZ |
| Ortiz-GilJ,2011 | Controls>Cognitivelypreservedgroup,Graymatter | 23 | F20 | Schizophrenia SZ |
| Paillere- | Schizophrenicpatients<Healthycontrols,Gray | 20 | F20 | Schizophrenia SZ |
| MartinotML,2001 |  |  |  |  |
| Pomarol-ClotetE,2010 | HealthyControls>SchizophreniaPatients,Gray | 31 | F20 | Schizophrenia SZ |
| Pomarol-ClotetE,2010 | HealthyControls>SchizophreniaPatients,Gray | 31 | F20 | Schizophrenia SZ |
| PriceG,2010 | SchizophreniaPatiets<HealthyControl,Greymatter | 47 | F20 | Schizophrenia SZ |
| QiuL,2011 | Schizophrenics<Controls,Graymatterdensity | 29 | F20 | Schizophrenia SZ |
| Salgado-PinedaP,2003 | Normals>SchizophreniaPatients,GrayMatter | 13 | F20 | Schizophrenia SZ |
| Salgado-PinedaP,2004 | Normals>SchizophreniaPatients,GrayMatter | 14 | F20 | Schizophrenia SZ |
| Salgado-PinedaP,2011 | Schizophrenicpatientsvs.Healthycontrols,Grey | 14 | F20 | Schizophrenia SZ |
| SchifferB,2013 | SZ+CD<HealthyControls,GrayMatterVolume | 25 | F20 | Schizophrenia SZ |
| SchifferB,2013 | SZ-CD<HealthyControls | 23 | F20 | Schizophrenia SZ |
| SchusterC,2012 | SchizophreniaPatients<HealthyControls,Greymatter | 27 | F20 | Schizophrenia SZ |
| ShapleskeJ,2002 | SchizophreniaPatients<Controls,GreyMatterVolume | 31 | F20 | Schizophrenia SZ |
| ShapleskeJ,2002 | Hallucinators<Controls,GreyMatterVolume | 32 | F20 | Schizophrenia SZ |
| ShapleskeJ,2002 | Non-hallucinators<Controls,GreyMatterVolume | 31 | F20 | Schizophrenia SZ |
| SigmundssonT,2001 | Schizophrenicpatientswithnegativesymptomsvs | 27 | F20 | Schizophrenia SZ |
| SmesnyS,2010 | HealthyControls>FirstEpisodePatients,GreyMatter | 13 | F20 | Schizophrenia SZ |
| SmesnyS,2010 | HealthyControls>RecurrentEpisodePatients,GreyMatter | 11 | F20 | Schizophrenia SZ |
| SuzukiM,2002 | Schizophreniapatients<HealthyControls,GrayMatter | 42 | F20 | Schizophrenia SZ |
| SuzukiM,2002 | Maleschizophreniapatients<Malehealthycontrols,Gray | 22 | F20 | Schizophrenia SZ |
| SuzukiM,2002 | Femaleschizophreniapatients<Femalehealthycontrols, | 20 | F20 | Schizophrenia SZ |
| ThebergeJ,2007 | HPAR2>30M,significantgreymattervolume | 16 | F20 | Schizophrenia SZ |
| TianL,2011 | Schizophrenicpatients<HC1,GrayMatterDensity | 30 | F20 | Schizophrenia SZ |
| TomelleriL,2009 | SchizophreniaPatients<HealthyControls,GrayMatter | 25 | F20 | Schizophrenia SZ |
| TregallasJR,2007 | HealthyControls>Schizophrenia,GrayMatterVolume | 32 | F20 | Schizophrenia SZ |
| Venkatasubramanian G,2008 | GMRegionsNegativelyCorrelatedwithMotor | 30 | F20 | Schizophrenia SZ |
| Venkatasubramanian G,2008 | HealthyControls>SchizophreniaPatients,Gray | 27 | F20 | Schizophrenia SZ |
| VoetsNL,2008 | CorrespondingGreyMatterDensityReductionsandSBM-based | 25 | F20 | Schizophrenia SZ |
| VoetsNL,2008 | GreyMatterDensityReductionsinPatientsvs.Controls | 25 | F20 | Schizophrenia SZ |
| WatsonDR,2012 | SZ<SCS,GrayMatterVolume | 25 | F20 | Schizophrenia SZ |
| WhitfordTJ,2006 | Normals>SchizophreniaPatientsatBaseline,Gray | 41 | F20 | Schizophrenia SZ |
| WhitfordTJ,2006 | Normals>SchizophreniaPatients,Session1>Session | 25 | F20 | Schizophrenia SZ |
| WilkeM,2001 | Normals>SchizophreniaPatients,GrayMatterVolume | 48 | F20 | Schizophrenia SZ |
| WolfRC,2008 | HealthyControls>Schizophrenia,GrayMatterConcentration | 14 | F20 | Schizophrenia SZ |
| WolfRC,2008 | HealthyControls>Schizophrenia,WhiteMatterVolume | 14 | F20 | Schizophrenia SZ |
| XuL,2009 | Normals>SchizophreniaPatients,GrayMatterConcentration | 120 | F20 | Schizophrenia SZ |
| YamadaM,2007 | SchizophreniaPatients<HealthyControls,GrayMatter | 20 | F20 | Schizophrenia SZ |
| YoshiharaY,2008 | HealthyControls>Schizophrenia,GreyMatter | 18 | F20 | Schizophrenia SZ |
| JanssenJ,2008 | PatientswithOtherPsychoses<HealthyControls,Gray | 25 | F28 | Other psychotic disorder not due to a substance or known physiological condition |
| KubickiM,2002 | Normals>AffectivePsychosisPatients,GrayMatter | 16 | F28 | Other psychotic disorder not due to a substance or known physiological condition |
| MarcelisM,2003 | Normals>Psychosis,GreyMatterDensity | 27 | F28 | Other psychotic disorder not due to a substance or known physiological condition |
| AdlemanNE,2012 | HV>BD,GrayMatterVolume | 55 | F31 | Bipolar Disorder BD, BPD |
| AdlerCM,2005 | Normals>BipolarDisorderPatients,GrayMatterVolume | 27 | F31 | Bipolar Disorder BD, BPD |
| AlmeidaJRC,2009 | BipolarDisordervsHealthyControl,GrayMatter | 27 | F31 | Bipolar Disorder BD, BPD |
| AlmeidaJRC,2009 | HealthyControlsvsBipolarDisorder,Non-apriori, | 27 | F31 | Bipolar Disorder BD, BPD |
| Alonso-LanaS,2016 | Cognitivelypreservedpatients<Controls,Greymatter | 28 | F31 | Bipolar Disorder BD, BPD |
| AmbrosiE,2013 | HealthyControls>BipolarPatients,GrayMatterVolume | 20 | F31 | Bipolar Disorder BD, BPD |
| CaiY,2015 | BipolarIdisorder<Healthycontrols,graymatter | 23 | F31 | Bipolar Disorder BD, BPD |
| ChenX,2007 | BDPatientswithFamilyHistoryvs.Normals,GrayMatter | 24 | F31 | Bipolar Disorder BD, BPD |
| ChenX,2007 | BDPatientswithoutFamilyHistoryvs.Normals,GrayMatter | 24 | F31 | Bipolar Disorder BD, BPD |
| ChenX,2007 | BDPatientswithPsychosisSymptomsvs.Normals,GrayMatter | 24 | F31 | Bipolar Disorder BD, BPD |
| ChenX,2007 | BDPatientswithoutPsychosisSymptomsvs.Normals,Gray | 24 | F31 | Bipolar Disorder BD, BPD |
| ChenX,2007 | BDPatientsTakingLithiumvs.Normals,GrayMatterDecreases | 24 | F31 | Bipolar Disorder BD, BPD |
| ChenX,2007 | BDPatientsNotTakingLithiumvs.Normals,GrayMatter | 24 | F31 | Bipolar Disorder BD, BPD |
| CuiL,2011 | BipolarManiaPatients<HealthyControls,GrayMatterVolume | 24 | F31 | Bipolar Disorder BD, BPD |
| DicksteinDP,2005 | Normals>PediatricBDPatients,GrayMatterVolume | 20 | F31 | Bipolar Disorder BD, BPD |
| DorisA,2004 | BD<CS,GrayMatterDensity | 11 | F31 | Bipolar Disorder BD, BPD |
| FarrowTFD,2005 | HealthyControls-InitialBipolarPatients,Reduced | 8 | F31 | Bipolar Disorder BD, BPD |
| GaoW,2013 | PediatricBipolarDisorder(PBD)<HC,Graymattervolume | 18 | F31 | Bipolar Disorder BD, BPD |
| HaldaneM,2008 | BD<CS,GrayMatterDensity | 44 | F31 | Bipolar Disorder BD, BPD |
| HallerS,2011 | Bipolarpatients<Healthycontrols,graymatter | 19 | F31 | Bipolar Disorder BD, BPD |
| HaTH,2010 | BipolarDisorderIIPatients<HealthyControls,GrayMatter | 23 | F31 | Bipolar Disorder BD, BPD |
| HaTH,2010 | BipolarDisorderIIPatients<HealthyControls,GrayMatter | 23 | F31 | Bipolar Disorder BD, BPD |
| JanssenJ,2008 | BipolarPatients<HealthyControls,GrayMatter | 20 | F31 | Bipolar Disorder BD, BPD |
| KimD,2013 | Bipolarpatients<Healthycontrols,graymatter | 49 | F31 | Bipolar Disorder BD, BPD |
| LadoucerCD,2008 | CONT>HBO,GrayMatterVolume | 20 | F31 | Bipolar Disorder BD, BPD |
| LiM,2011 | BD<CS,GrayMatterVolume | 24 | F31 | Bipolar Disorder BD, BPD |
| LyooIK,2004 | Normals>BipolarDiseasePatients,GrayMatterDensity | 39 | F31 | Bipolar Disorder BD, BPD |
| McIntoshAM,2004 | Normals>BDPatientsfromMixedFamilies,GrayMatter | 19 | F31 | Bipolar Disorder BD, BPD |
| MolinaV,2011 | BD<CS,GrayMatterVolume | 19 | F31 | Bipolar Disorder BD, BPD |
| NaritaK,2011 | BipolarIIPatientswithrapidcycling<Healthycontrols, | 14 | F31 | Bipolar Disorder BD, BPD |
| NaritaK,2011 | BipolarIIPatientswithoutrapidcycling<Healthy | 17 | F31 | Bipolar Disorder BD, BPD |
| NugentAC,2006 | Normals>UnmedicatedBipolarDisorder(BD)Patients, | 16 | F31 | Bipolar Disorder BD, BPD |
| NugentAC,2006 | Normals>BDMedicated,ApproachingSignificance,Gray | 20 | F31 | Bipolar Disorder BD, BPD |
| RedlichR,2014 | Bipolarpatients<Healthycontrols,graymatter | 58 | F31 | Bipolar Disorder BD, BPD |
| RossiR,2012 | BipolarDisorder(BD)<HC,graymattervolume | 14 | F31 | Bipolar Disorder BD, BPD |
| SaricicekA,2015 | BipolarIpatients<healthycontrols,graymatter | 28 | F31 | Bipolar Disorder BD, BPD |
| SinghMK,2012 | BipolarIDisorderPatients(BDI)<HC,Graymatter | 24 | F31 | Bipolar Disorder BD, BPD |
| SinghMK,2012 | BDI<HC,Graymattervolume,p<0.001 | 24 | F31 | Bipolar Disorder BD, BPD |
| StanfieldAC,2009 | BipolarDisorderPatients<HealthyControls,Gray | 66 | F31 | Bipolar Disorder BD, BPD |
| TangLR,2014 | Healthycontrols>bipolarIpatients,graymatter | 27 | F31 | Bipolar Disorder BD, BPD |
| TostH,2010 | HealthySubjects>(BD)BipolarDisorderPatients | 42 | F31 | Bipolar Disorder BD, BPD |
| TostH,2010 | HealthySubjects>BipolarDisorderPatientswithPsychotic | 30 | F31 | Bipolar Disorder BD, BPD |
| TostH,2010 | HealthtySubjects>BipolarDisorderPatientswithPersecutory | 15 | F31 | Bipolar Disorder BD, BPD |
| WangF,2011 | BD<CS,GrayMatterVolume | 41 | F31 | Bipolar Disorder BD, BPD |
| WatsonDR,2012 | BD<bCS,GrayMatterVolume | 24 | F31 | Bipolar Disorder BD, BPD |
| AbeO,2010 | UnipolarDepressiveDisorderPatients<HealthyControls,Gray | 21 | F32-F33 | Major Depressive Disorder |
| AlemanyS,2013 | UnaffectedTwin>AffectedTwin,GrayMatterVolume | 10 | F32-F33 | Major Depressive Disorder |
| AlemanyS,2013 | HC>CTMDDGrayMatterVolume | 12 | F32-F33 | Major Depressive Disorder |
| ArnoneD,2009 | Depression<Controls,GrayMatterVolume | 25 | F32-F33 | Major Depressive Disorder |
| ArnoneD,2013 | Currentlydepressedpatients(cMDD)<Healthycontrols(HC) | 39 | F32-F33 | Major Depressive Disorder |
| BergouignanL,2009 | UnipolarDepressedPatients<HealthyControls,Grey | 21 | F32-F33 | Major Depressive Disorder |
| CaiY,2015 | MDDpatients<Healthycontrols,graymatter | 23 | F32-F33 | Major Depressive Disorder |
| ChaneyA,2014 | Control<Majordepressivedisorderpatients(MDD) | 10 | F32-F33 | Major Depressive Disorder |
| ChengY,2010 | MDDPatients<Controls,GrayMatterVolume | 68 | F32-F33 | Major Depressive Disorder |
| ChengY,2010 | HDRSScorevs.GrayMatterVolume,PositiveCorrelation | 68 | F32-F33 | Major Depressive Disorder |
| EggerK,2008 | HealthyControls>DepressivePatients,GrayMatterVolume | 14 | F32-F33 | Major Depressive Disorder |
| FrodlT,2008 | MDDPatients<Controls,GrayMatterDensity | 30 | F32-F33 | Major Depressive Disorder |
| GongQ,2011 | HealthyControls(HC)>RefractoryDepressiveDisorder(RDD) | 23 | F32-F33 | Major Depressive Disorder |
| GongQ,2011 | HC>Non-RefractoryDepressiveDisorder(NDD) | 23 | F32-F33 | Major Depressive Disorder |
| GrieveSM,2013 | Controls>MDD,Corticalthickness | 34 | F32-F33 | Major Depressive Disorder |
| GrieveSM,2013 | Controls>MajorDepressiveDisorder(MDD) | 34 | F32-F33 | Major Depressive Disorder |
| GuoW,2014 | Majordepressivedisorderpatients(MDD)<Healthycontrols | 44 | F32-F33 | Major Depressive Disorder |
| GuoW,2014 | Firstepisodepatients<controls | 24 | F32-F33 | Major Depressive Disorder |
| GuoW,2014 | Recurrentpatients<controls | 21 | F32-F33 | Major Depressive Disorder |
| HwangJ,2010 | DepressionPatients<HealthyControls,GrayMatterVolume | 26 | F32-F33 | Major Depressive Disorder |
| InksterB,2011 | AllMDDPatientsvs.HealthyControls,GrayMatterVolume | 49 | F32-F33 | Major Depressive Disorder |
| KimMJ,2008 | HealthyControls>MajorDepressiveDisorderPatients,Gray | 22 | F32-F33 | Major Depressive Disorder |
| LaiCH,2015 | HealthyControls>MajorDepressiveDisorder(MDD)Patients, | 53 | F32-F33 | Major Depressive Disorder |
| LeeHY,2011 | HealthyControls>MDDPatients,GrayMatterConcentration | 47 | F32-F33 | Major Depressive Disorder |
| LeungKK,2009 | HealthyControls>DepressedPatients,GrayMatter | 17 | F32-F33 | Major Depressive Disorder |
| LeungKK,2009 | HealthyControls>DepressedPatients,GrayMatterVolume | 17 | F32-F33 | Major Depressive Disorder |
| LiCT,2010 | Non-RemitingDepressives<HealthyControls,GrayMatterVolume | 25 | F32-F33 | Major Depressive Disorder |
| LiCT,2010 | RemittingMDDPatients<HealthyControls,GrayMatterVolume | 19 | F32-F33 | Major Depressive Disorder |
| LiuCH,2014 | CurrentdiagnosisofMajorDepressiveDisorder(cMDD)< | 19 | F32-F33 | Major Depressive Disorder |
| LiuCH,2014 | PrevioushistoryofdiagnosableMajorDepressiveDisorder | 19 | F32-F33 | Major Depressive Disorder |
| MakAK,2009 | MajorDepressiveDisorderPatients<HealthyControls,Gray | 17 | F32-F33 | Major Depressive Disorder |
| MakAK,2009 | MajorDepressiveDisorderPatients<HealthyControls,Gray | 17 | F32-F33 | Major Depressive Disorder |
| PengJ,2010 | MajorDepressiveDisorderPatients<HealthyControls,Gray | 22 | F32-F33 | Major Depressive Disorder |
| RedlichR,2014 | MDDpatients<Healthycontrols,graymatter | 58 | F32-F33 | Major Depressive Disorder |
| RiesML,2009 | HealthyControls>DepressedGroup,GrayMatterVolume | 15 | F32-F33 | Major Depressive Disorder |
| SalvadoreG,2011 | ChronicMDDPatients<HealthyControls,GrayMatter | 58 | F32-F33 | Major Depressive Disorder |
| SalvadoreG,2011 | RemissionMDDPatients<HealthyControls,GrayMatter | 27 | F32-F33 | Major Depressive Disorder |
| SalvadoreG,2011 | ChronicMDDPatients<HealthyControls,GrayMatter | 58 | F32-F33 | Major Depressive Disorder |
| SalvadoreG,2011 | RemissionMDDPatients<HealthyControls,GrayMatter | 27 | F32-F33 | Major Depressive Disorder |
| ScheuereckerJ,2010 | MDDPatients<HealthyControls,GrayMatterVolume | 13 | F32-F33 | Major Depressive Disorder |
| Serra-BlascoM,2013 | HC>tMDD,GrayMatterVolume | 22 | F32-F33 | Major Depressive Disorder |
| Serra-BlascoM,2013 | HC>rMDD,GrayMatterVolume | 22 | F32-F33 | Major Depressive Disorder |
| ShadMU,2012 | HC>Dep,GrayMatterVolume | 22 | F32-F33 | Major Depressive Disorder |
| ShahPJ,1998 | Treatment-Resistant<HealthyControls,GreyMatterDensity | 20 | F32-F33 | Major Depressive Disorder |
| ShahPJ,1998 | HealthyControls<Treatment-Resistant,GreyMatterDensity | 20 | F32-F33 | Major Depressive Disorder |
| ShahPJ,1998 | GreyMatterDensityvs.DelayedVerbalRecognition | 20 | F32-F33 | Major Depressive Disorder |
| Soriano-MasC,2011 | OriginalMDDPatients<OriginalHealthyControls,Gray | 40 | F32-F33 | Major Depressive Disorder |
| StratmannM,2014 | Controls>Patients | 132 | F32-F33 | Major Depressive Disorder |
| StratmannM,2014 | Control>Patientswithrecurrentdepressivedisorder | 97 | F32-F33 | Major Depressive Disorder |
| TakiY,2005 | Normals>Geriatricsubthresholddepressionpatients,gray | 13 | F32-F33 | Major Depressive Disorder |
| vanEijndhovenP,2013 | CorticalThickness,MDDPatients<HealthyControls | 20 | F32-F33 | Major Depressive Disorder |
| vanTolMJ,2010 | MDD<HealthyControls,GrayMatterVolume | 65 | F32-F33 | Major Depressive Disorder |
| vanTolMJ,2013 | MajorDepressiveDisorder>Healthycontrols,Lower | 20 | F32-F33 | Major Depressive Disorder |
| WagnerG,2008 | HealthyControls>MDDPatients,GreyMatterVolume | 15 | F32-F33 | Major Depressive Disorder |
| WagnerG,2011 | MDDPatients<HealthyControls,GrayMatterVolume | 30 | F32-F33 | Major Depressive Disorder |
| WagnerG,2011 | HighRiskforSuicideMDDPatients<HealthyControls,Gray | 15 | F32-F33 | Major Depressive Disorder |
| ZhangT,2009 | TRD<Controls,GrayMatterVolume(Voxel-Wise) | 15 | F32-F33 | Major Depressive Disorder |
| ZhangT,2009 | TRD<Controls,GrayMatterVolume(Cluster-Wise) | 15 | F32-F33 | Major Depressive Disorder |
| ZhangX,2012 | HC>CVD,GrayMatterVolume | 30 | F32-F33 | Major Depressive Disorder |
| ZhangX,2012 | HC>MDD,GrayMatterVolume | 32 | F32-F33 | Major Depressive Disorder |
| ZhangX,2012 | MDD>HC,GrayMatterVolume | 32 | F32-F33 | Major Depressive Disorder |
| ZhangX,2012 | GrayMatterVolume,MainEffectofGroup | 30 | F32-F33 | Major Depressive Disorder |
| ZouK,2010 | Depression<Controls,GrayMatterVolume | 23 | F32-F33 | Major Depressive Disorder |
| GilbertAR,2008 | HealthyControls>OCDPatients,GrayMatterVolume | 20 | F42 | Obsessive Compulsive Disorder OCD |
| KoprivovaJ,2009 | ObsessiveCompulsiveDisorderPatients<Healthy | 14 | F42 | Obsessive Compulsive Disorder OCD |
| MatsumotoR,2010 | OCD<CS,GrayMatterVolume | 16 | F42 | Obsessive Compulsive Disorder OCD |
| PujolJ,2004 | OCDPatients<HealthyControls,absoluteGreyMattervolume | 72 | F42 | Obsessive Compulsive Disorder OCD |
| SzeszkoPR,2008 | HealthyControls>ObsessiveCompulsiveDisorder | 26 | F42 | Obsessive Compulsive Disorder OCD |
| TogaoO,2010 | OCDpatients<Controls,GrayMatterVolume | 23 | F42 | Obsessive Compulsive Disorder OCD |
| ValenteAAJr,2005 | Normals>OverallOCDPatients,GrayMatterVolume | 15 | F42 | Obsessive Compulsive Disorder OCD |
| ValenteAAJr,2005 | Normals>OCDPatientsWithoutMajorDepression,Gray | 15 | F42 | Obsessive Compulsive Disorder OCD |
| vandenHeuvelOA,2009 | Obsessive-CompulsiveDisorderPatients<Healthy | 50 | F42 | Obsessive Compulsive Disorder OCD |
| vandenHeuvelOA,2009 | NegativeCorrelationswiththeHarmChecking | 55 | F42 | Obsessive Compulsive Disorder OCD |
| YooSY,2008 | FemaleOCDPatients<FemaleHealthyControls,GrayMatter | 24 | F42 | Obsessive Compulsive Disorder OCD |
| YooSY,2008 | MaleOCDPatients<MaleHealthyControls,GrayMatter | 47 | F42 | Obsessive Compulsive Disorder OCD |
| LuC,2010 | HealthyControls>StutteringSpeakers,GrayMatterVolume | 12 | F80 | Specific Developmental Disorders of Speech and Language |
| WatkinsKE,2002 | UnrelatedNormals>AffectedSpeechandLanguage | 10 | F80 | Specific Developmental Disorders of Speech and Language |
| AbellF,1999 | AutisticDisorderPatients<HealthyControls,GrayMatter | 15 | F84 | Pervasive Developmental Disorders PDD, autism, ASD |
| BrieberS,2007 | Controls>ASD,GreyMatterDifferences | 15 | F84 | Pervasive Developmental Disorders PDD, autism, ASD |
| ChengY,2011 | HealthyControls>AutismSpectrumDisorderPatients,Grey | 25 | F84 | Pervasive Developmental Disorders PDD, autism, ASD |
| ChengY,2011 | HealthyControls>Asperger'sSyndromePatients,GrayMatter | 11 | F84 | Pervasive Developmental Disorders PDD, autism, ASD |
| ChengY,2011 | HealthyControls>AutismPatients,GrayMatterVolume | 12 | F84 | Pervasive Developmental Disorders PDD, autism, ASD |
| ChengY,2011 | HealthyControls>AutismSpectrumDisorderPatients,Gray | 25 | F84 | Pervasive Developmental Disorders PDD, autism, ASD |
| ChengY,2011 | HealthyControls>Asperger'sSyndromePatients,GrayMatter | 11 | F84 | Pervasive Developmental Disorders PDD, autism, ASD |
| ChengY,2011 | HealthyControls>AutismPatients,GrayMatter | 12 | F84 | Pervasive Developmental Disorders PDD, autism, ASD |
| CraigMC,2007 | Autism-SpectrumDisorderPatients<HealthyControls,Gray | 14 | F84 | Pervasive Developmental Disorders PDD, autism, ASD |
| EckerC,2010 | ASD<Controls,Graymattervolume | 22 | F84 | Pervasive Developmental Disorders PDD, autism, ASD |
| EckerC,2012 | HealthyControls>ASDPatients,GrayMatterVolume | 89 | F84 | Pervasive Developmental Disorders PDD, autism, ASD |
| EckerC,2012 | HealthyControls>ASDPatients,GrayMatterVolume | 89 | F84 | Pervasive Developmental Disorders PDD, autism, ASD |
| HydeKL,2010 | HealthyControls>AutismPatients,GrayMatter | 13 | F84 | Pervasive Developmental Disorders PDD, autism, ASD |
| KeX,2008 | HealthyControls>High-FunctioningAutismPatients,Gray | 15 | F84 | Pervasive Developmental Disorders PDD, autism, ASD |
| KosakaH,2010 | HealthyControls>PervasiveDevelopmentalDisorder,Gray | 32 | F84 | Pervasive Developmental Disorders PDD, autism, ASD |
| KurthF,2011 | AutismPatients<HealthyControls,GrayMatterVolume | 52 | F84 | Pervasive Developmental Disorders PDD, autism, ASD |
| KwonH,2004 | HealthyControls-AspergerSyndromePatients,GrayMatter | 11 | F84 | Pervasive Developmental Disorders PDD, autism, ASD |
| KwonH,2004 | HealthyControls-PervasiveDevelopmentalDisorder(PDD) | 9 | F84 | Pervasive Developmental Disorders PDD, autism, ASD |
| McAlonanGM,2002 | Asperger'sSyndromePatients<HealthyControls,Gray | 17 | F84 | Pervasive Developmental Disorders PDD, autism, ASD |
| McAlonanGM,2005 | AutismPatients<HealthyControls,GrayMatterVolume | 17 | F84 | Pervasive Developmental Disorders PDD, autism, ASD |
| McAlonanGM,2008 | High-FunctioningAutismPatients<HealthyControls, | 17 | F84 | Pervasive Developmental Disorders PDD, autism, ASD |
| McAlonanGM,2008 | Asperger'sSyndromePatients<HealthyControls,Gray | 16 | F84 | Pervasive Developmental Disorders PDD, autism, ASD |
| MengottiP,2011 | HealthyControls>AutismPatients,GrayMatterVolume | 20 | F84 | Pervasive Developmental Disorders PDD, autism, ASD |
| RivaD,2011 | AutismSpectrumDisorderPatients<HealthyControls,Gray | 21 | F84 | Pervasive Developmental Disorders PDD, autism, ASD |
| SalmondCH,2007 | AutismSpectrumDisorderPatients<HealthyControls, | 9 | F84 | Pervasive Developmental Disorders PDD, autism, ASD |
| SpencerMD,2006 | Normals>IntellectuallyDisabledPatients,GreyMatter | 15 | F84 | Pervasive Developmental Disorders PDD, autism, ASD |
| ToalF,2010 | AutismSpectrumDisorderPatients<HealthyControls,Gray | 33 | F84 | Pervasive Developmental Disorders PDD, autism, ASD |
| ToalF,2010 | AspergerSyndromePatientSubgroup<HealthyControls,Gray | 33 | F84 | Pervasive Developmental Disorders PDD, autism, ASD |
| ToalF,2010 | AutismPatientSubgroup<HealthyControls,GrayMatter | 26 | F84 | Pervasive Developmental Disorders PDD, autism, ASD |
| AhrendtsJ,2011 | ADHD<CS,GrayMatterVolume | 31 | F90 | Attention Deficit/Hyperactivity Disorder |
| BrieberS,2007 | Controls>ADHD,GreyMatterDifferences | 15 | F90 | Attention Deficit/Hyperactivity Disorder |
| CarmonaS,2005 | HealthyControls>ADHDPatients,GrayMatterVolume | 25 | F90 | Attention Deficit/Hyperactivity Disorder |
| KobelM,2010 | ADHD<HealthyControls,GrayMatter | 12 | F90 | Attention Deficit/Hyperactivity Disorder |
| McAlonanGM,2007 | HealthyControls>ADHDPatients,GrayMatterVolume | 10 | F90 | Attention Deficit/Hyperactivity Disorder |
| OvermeyerS,2001 | ADHD<HealthyControls,GrayMatterVolume | 16 | F90 | Attention Deficit/Hyperactivity Disorder |
| SasayamaD,2010 | ADHD+ODDCD<HC,GrayMatterDensity | 10 | F90 | Attention Deficit/Hyperactivity Disorder |
| SasayamaD,2010 | ADHD+(ADHD+ODDCD)<HC,GrayMatterVolume | 8 | F90 | Attention Deficit/Hyperactivity Disorder |
| WangJ,2007 | ADHD<CS,GrayMatterDensity | 12 | F90 | Attention Deficit/Hyperactivity Disorder |
| LudolphAG,2006 | Normals>TouretteSyndromeAdolescents,GreyMatter | 14 | F95 | Other Disorders of Psychological Development. Tourette |
| Muller-VahlKR,2009 | TouretteSyndromePatients>Controls,GrayMatter | 19 | F95 | Other Disorders of Psychological Development. Tourette |

**Table S2:** experiments included in the control meta-analysis of increase-only co-alteration.

| AUTHOR | CONDITION | # SUBJECTS | ICD-10 CODE | DIAGNOSIS |
| --- | --- | --- | --- | --- |
| AntonovaE,2005 | SchizophreniaPatients>Normals,GreyMatterVolume | 40 | F20 | Schizophrenia SZ |
| BassittDP,2007 | Schizophrenia>Normals,GrayMatterVolume | 30 | F20 | Schizophrenia SZ |
| BrownGG,2011 | Healthycontrols<Schizophreniapatients,graymatter | 17 | F20 | Schizophrenia SZ |
| CalhounVD,2006 | SchizophreniaPatients>Normals,GrayMatterVolume | 15 | F20 | Schizophrenia SZ |
| CuiL,2011 | Paranoid-typeSchizophreniaPatients>HealthyControls,Gray | 23 | F20 | Schizophrenia SZ |
| DengMY,2009 | Patients[re-scanwithin3weeks]>Healthycontrolssecond | 10 | F20 | Schizophrenia SZ |
| DengMY,2009 | Patients[re-scanbeyond3weeks]>Healthycontrols, | 10 | F20 | Schizophrenia SZ |
| GiulianiNR,2005 | SchizophreniaPatients>HealthyControls,GrayMatter | 34 | F20 | Schizophrenia SZ |
| HaTH,2004 | ParanoidSchizophreniaPatients>HealthyControls,Gray | 35 | F20 | Schizophrenia SZ |
| HoneaRA,2008 | SchizophreniaSpectrumDisorder>HealthyControls,Gray | 169 | F20 | Schizophrenia SZ |
| HulshoffPolHE,2001 | SchizophreniaPatients>Normals,GrayMatter | 158 | F20 | Schizophrenia SZ |
| HydeTM,2008 | SchizoprheniaPatientswithEnuresis<Schizophrenia | 16 | F20 | Schizophrenia SZ |
| KasparekT,2010 | First-EpisodeSchizophrenia>HealthyControls,Gray | 49 | F20 | Schizophrenia SZ |
| KawasakiY,2004 | SchizophreniaPatients>HealthyControls,GrayMatter | 25 | F20 | Schizophrenia SZ |
| McDonaldC,2005 | Schizophrenia>Normals,GrayMatterVolume | 25 | F20 | Schizophrenia SZ |
| MolinaV,2011 | SZ>CS,GrayMatterVolume | 24 | F20 | Schizophrenia SZ |
| MolinaV,2011 | Patients>Healthycontrols,increasesingreymattervolume | 30 | F20 | Schizophrenia SZ |
| MoriyaJ,2010 | SchizophreniaPatients>HealthyControls(DiffusionTensor | 19 | F20 | Schizophrenia SZ |
| O'DalyO,2007 | SchizophreniaPatients>HealthyControls,Greymatter | 28 | F20 | Schizophrenia SZ |
| PriceG,2010 | SchizophreniaPatiets>HealthyControl,Greymatter | 47 | F20 | Schizophrenia SZ |
| Salgado-PinedaP,2003 | SchizophreniaPatients>Normals,GrayMatter | 13 | F20 | Schizophrenia SZ |
| SchifferB,2013 | SZ+CD>HealthyControls,GrayMatterVolume | 25 | F20 | Schizophrenia SZ |
| ShapleskeJ,2002 | SchizophreniaPatients>Controls,GreyMatterVolume | 31 | F20 | Schizophrenia SZ |
| ShapleskeJ,2002 | Hallucinators>Controls,GreyMatterVolume | 32 | F20 | Schizophrenia SZ |
| SmesnyS,2010 | FirstEpisodePatients>HealthyControls,GreyMatter | 13 | F20 | Schizophrenia SZ |
| SmesnyS,2010 | RecurrentEpisodePatients>HealthyControls,GreyMatter | 11 | F20 | Schizophrenia SZ |
| SuzukiM,2002 | Schizophreniapatients>HealthyControls,GrayMatter | 42 | F20 | Schizophrenia SZ |
| SuzukiM,2002 | Femaleschizophreniapatients>Femalehealthycontrols, | 20 | F20 | Schizophrenia SZ |
| TanskanenP,2010 | Schizophrenia<Controls,GrayMatterDensity | 54 | F20 | Schizophrenia SZ |
| ThebergeJ,2007 | NT>30M,significantgreymattervolume | 16 | F20 | Schizophrenia SZ |
| WatsonDR,2012 | SZ>sCS,GrayMatterVolume | 25 | F20 | Schizophrenia SZ |
| WhitfordTJ,2006 | SchizophreniaPatients>NormalsatBaseline,Gray | 41 | F20 | Schizophrenia SZ |
| WilkeM,2001 | SchizophreniaPatients>Normals,GrayMatterVolume | 48 | F20 | Schizophrenia SZ |
| deCastro-ManganoP,2011 | AreasofGreaterLongitudinalGrayMatter | 17 | F28 | Other psychotic disorder not due to a substance or known physiological condition |
| MarcelisM,2003 | Psychosis>Normals,GreyMatterDensity | 27 | F28 | Other psychotic disorder not due to a substance or known physiological condition |
| MarcelisM,2003 | Psychosis>Relatives,GrayMatterDensity | 31 | F28 | Other psychotic disorder not due to a substance or known physiological condition |
| AdlerCM,2005 | BipolarDisorderPatients>Normals,GrayMatterVolume | 27 | F31 | Bipolar Disorder BD, BPD |
| AdlerCM,2007 | BipolarDisorderPatients>Normals,GrayMatterVolume | 33 | F31 | Bipolar Disorder BD, BPD |
| AdlerCM,2007 | BipolarDisorderPatients>Normals,GrayMatterDensity | 33 | F31 | Bipolar Disorder BD, BPD |
| BrownGG,2011 | Healthycontrols<Bipolarpatients,graymatter | 15 | F31 | Bipolar Disorder BD, BPD |
| ChenX,2007 | BDPatientswithFamilyHistoryvs.Normals,GrayMatter | 24 | F31 | Bipolar Disorder BD, BPD |
| ChenX,2007 | BDPatientswithoutFamilyHistoryvs.Normals,GrayMatter | 24 | F31 | Bipolar Disorder BD, BPD |
| ChenX,2007 | BDPatientswithPsychosisSymptomsvs.Normals,GrayMatter | 24 | F31 | Bipolar Disorder BD, BPD |
| ChenX,2007 | BDPatientsNotTakingLithiumvs.Normals,GrayMatter | 24 | F31 | Bipolar Disorder BD, BPD |
| ChenZ,2012 | Bipolarpatients>Healthycontrols,greymattervolume | 18 | F31 | Bipolar Disorder BD, BPD |
| CuiL,2011 | BipolarManiaPatients>HealthyControls,GrayMatterVolume | 24 | F31 | Bipolar Disorder BD, BPD |
| FrangouS,2012 | Bipolarpatients>Healthycontrols,Greymattervolume | 47 | F31 | Bipolar Disorder BD, BPD |
| HaldaneM,2008 | BD>CS,GrayMatterDensity | 44 | F31 | Bipolar Disorder BD, BPD |
| HaTH,2010 | BipolarDisorderIPatients<BipolarDisorderIIPatients, | 23 | F31 | Bipolar Disorder BD, BPD |
| KemptonMJ,2009 | MainEffectofGroupinBipolarDisorderPatients, | 30 | F31 | Bipolar Disorder BD, BPD |
| KozickyJM,2013 | BipolarIpatients>Healthycontrols,graymatter | 30 | F31 | Bipolar Disorder BD, BPD |
| LadoucerCD,2008 | HBO>CONT,GrayMatterVolume | 20 | F31 | Bipolar Disorder BD, BPD |
| PericoCAM,2011 | BD>HC,GrayMatterVolume | 26 | F31 | Bipolar Disorder BD, BPD |
| SaricicekA,2015 | BipolarIpatients>Healthycontrols,graymatter | 28 | F31 | Bipolar Disorder BD, BPD |
| TangLR,2014 | Helathycontrols<bipolarIpatients,graymatter | 27 | F31 | Bipolar Disorder BD, BPD |
| WatsonDR,2012 | BD>bCS,GrayMatterVolume | 24 | F31 | Bipolar Disorder BD, BPD |
| AmicoF,2011 | Controlswithfamilyhistorydepression(HC-FHP)<MDD | 30 | F32-F33 | Major Depressive Disorder |
| ArnoneD,2009 | Controls<Depression,GrayMatterVolume | 25 | F32-F33 | Major Depressive Disorder |
| ArnoneD,2013 | cMDD>Healthycontrols,longitudinalstudy | 15 | F32-F33 | Major Depressive Disorder |
| ArnoneD,2013 | rMDD>subgroupofhealthycontrols,longitudinalstudy | 15 | F32-F33 | Major Depressive Disorder |
| BiedermannSV,2015 | Age-DependentGrayMatterVolumeLoss,Patients> | 35 | F32-F33 | Major Depressive Disorder |
| ChaneyA,2014 | Control<MDD | 10 | F32-F33 | Major Depressive Disorder |
| ChaneyA,2014 | Controlnomaltreatment<MDDnomaltreatment | 17 | F32-F33 | Major Depressive Disorder |
| ChaneyA,2014 | Controlmaltreatment<MDDmaltreatment | 10 | F32-F33 | Major Depressive Disorder |
| GongQ,2011 | RDD>HC | 23 | F32-F33 | Major Depressive Disorder |
| GongQ,2011 | NDD>HC | 23 | F32-F33 | Major Depressive Disorder |
| GrieveSM,2013 | MDD>Controls,Corticalthickness | 34 | F32-F33 | Major Depressive Disorder |
| HwangJ,2010 | HealthyControls<DepressionPatients,GrayMatterVolume | 26 | F32-F33 | Major Depressive Disorder |
| LeungKK,2009 | DepressedPatients>HealthyControls,GrayMatterVolume | 17 | F32-F33 | Major Depressive Disorder |
| PericoCAM,2011 | HC>MDD,GrayMatterVolume | 20 | F32-F33 | Major Depressive Disorder |
| QiuL,2014 | MajorDepressedDisorderpatients(MDD)>Healthycontrols, | 46 | F32-F33 | Major Depressive Disorder |
| QiuL,2014 | MDD>Healthycontrols,GrayMatterVolume | 46 | F32-F33 | Major Depressive Disorder |
| ScheuereckerJ,2010 | HealthyControls<MDDPatients,GrayMatterVolume | 13 | F32-F33 | Major Depressive Disorder |
| TruongW,2013 | Early-onsetMajorDepresseddisorderpatients(MDD)> | 28 | F32-F33 | Major Depressive Disorder |
| vanEijndhovenP,2013 | CorticalThickness,HealthyControls<MDDPatients | 20 | F32-F33 | Major Depressive Disorder |
| ChristianCJ,2008 | OCDPatients>HealthyControls,GrayMatter | 21 | F42 | Obsessive Compulsive Disorder OCD |
| ChristianCJ,2008 | OCDPatientsWithoutMajorDepression>Healthy | 21 | F42 | Obsessive Compulsive Disorder OCD |
| GilbertAR,2008 | OCDPatients>HealthyControls,GrayMatterVolume | 20 | F42 | Obsessive Compulsive Disorder OCD |
| KimJJ,2001 | OCD>CS,GrayMatterVolume | 25 | F42 | Obsessive Compulsive Disorder OCD |
| KimJJ,2001 | OCD<CS,GrayMatterVolume | 25 | F42 | Obsessive Compulsive Disorder OCD |
| PujolJ,2004 | OCDPatients>HealthyControls,relativeGreyMattervolume | 72 | F42 | Obsessive Compulsive Disorder OCD |
| SzeszkoPR,2008 | ObsessiveCompulsiveDisorderPatients>Healthy | 26 | F42 | Obsessive Compulsive Disorder OCD |
| ValenteAAJr,2005 | OverallOCDPatients>Normals,GrayMatterVolume | 15 | F42 | Obsessive Compulsive Disorder OCD |
| ValenteAAJr,2005 | OCDPatientsWithoutMajorDepression>Normals,Gray | 15 | F42 | Obsessive Compulsive Disorder OCD |
| YooSY,2008 | MaleOCDPatients>MaleHealthyControls,GrayMatter | 47 | F42 | Obsessive Compulsive Disorder OCD |
| BealDS,2007 | StutteringPatients>HealthyControls,GrayMatterDensity | 26 | F80 | Specific Developmental Disorders of Speech and Language |
| LuC,2010 | StutteringSpeakers>HealthyControls,GrayMatterVolume | 12 | F80 | Specific Developmental Disorders of Speech and Language |
| WatkinsKE,2002 | AffectedSpeechandLanguageDisorderPatients> | 10 | F80 | Specific Developmental Disorders of Speech and Language |
| AbellF,1999 | AutisticDisorderPatients>HealthyControls,GrayMatter | 15 | F84 | Pervasive Developmental Disorders PDD, autism, ASD |
| BonilhaL,2008 | ClassicAutismPatients>HealthyControls,GrayMatter | 12 | F84 | Pervasive Developmental Disorders PDD, autism, ASD |
| BrieberS,2007 | ConjunctionAnalysis | 15 | F84 | Pervasive Developmental Disorders PDD, autism, ASD |
| BrieberS,2007 | ConjunctionAnalysis | 15 | F84 | Pervasive Developmental Disorders PDD, autism, ASD |
| BrieberS,2007 | ASD>Controls,GreyMatterDifferences | 15 | F84 | Pervasive Developmental Disorders PDD, autism, ASD |
| CalderoniS,2012 | FemaleAutismSpectrumDisordersPatients>Female | 19 | F84 | Pervasive Developmental Disorders PDD, autism, ASD |
| ChengY,2011 | AutismSpectrumDisorderPatients>HealthyControls,Grey | 25 | F84 | Pervasive Developmental Disorders PDD, autism, ASD |
| ChengY,2011 | Asperger'sSyndromePatients>HealthyControls,GrayMatter | 11 | F84 | Pervasive Developmental Disorders PDD, autism, ASD |
| ChengY,2011 | AutismPatients>HealthyControls,GrayMatterVolume | 12 | F84 | Pervasive Developmental Disorders PDD, autism, ASD |
| ChengY,2011 | AutismSpectrumDisorderPatients>HealthyControls,Gray | 25 | F84 | Pervasive Developmental Disorders PDD, autism, ASD |
| ChengY,2011 | Asperger'sSyndromePatients>HealthyControls,GrayMatter | 11 | F84 | Pervasive Developmental Disorders PDD, autism, ASD |
| ChengY,2011 | AutismPatients>HealthyControls,GrayMatter | 12 | F84 | Pervasive Developmental Disorders PDD, autism, ASD |
| EckerC,2010 | ASD>Controls,Graymattervolume | 22 | F84 | Pervasive Developmental Disorders PDD, autism, ASD |
| EckerC,2012 | ASDPatients>HealthyControls,GrayMatterVolume | 89 | F84 | Pervasive Developmental Disorders PDD, autism, ASD |
| EckerC,2012 | ASDPatients>HealthyControls,GrayMatterVolume | 89 | F84 | Pervasive Developmental Disorders PDD, autism, ASD |
| HendryJ,2006 | AutismPatients>Normals | 19 | F84 | Pervasive Developmental Disorders PDD, autism, ASD |
| HydeKL,2010 | AutismPatients>HealthyControls,GrayMatter | 13 | F84 | Pervasive Developmental Disorders<br>PDD, autism, ASD |
| KeX,2008 | High-FunctioningAutismPatients>HealthyControls,Gray | 15 | F84 | Pervasive Developmental Disorders<br>PDD, autism, ASD |
| MengottiP,2011 | AutismPatients>HealthyControls,GrayMatterVolume | 20 | F84 | Pervasive Developmental Disorders<br>PDD, autism, ASD |
| SalmondCH,2007 | AutismSpectrumDisorderPatients>HealthyControls, | 9 | F84 | Pervasive Developmental Disorders<br>PDD, autism, ASD |
| SchmitzN,2006 | AutisticSpectrumDisorder>HealthyControls,GrayMatter | 10 | F84 | Pervasive Developmental Disorders<br>PDD, autism, ASD |
| ToalF,2010 | AutismPatientSubgroup>HealthyControls,GrayMatter | 26 | F84 | Pervasive Developmental Disorders<br>PDD, autism, ASD |
| WaiterGD,2004 | AutismSpectrumDisorderPatients>HealthyControls, | 16 | F84 | Pervasive Developmental Disorders<br>PDD, autism, ASD |
| BrieberS,2007 | ADHD>Controls,GreyMatterDifferences | 15 | F90 | Attention Deficit/Hyperactivity Disorder |
| WangJ,2007 | ADHD>CS,GrayMatterDensity | 12 | F90 | Attention Deficit/Hyperactivity Disorder |
| GarrauxG,2006 | Tourette'sSyndrome>Normals,GrayMatterVolume | 31 | F95 | Other Disorders of Psychological<br>Development. Tourette |
| LudolphAG,2006 | TouretteSyndromeAdolescents>Normals,GreyMatter | 14 | F95 | Other Disorders of Psychological<br>Development. Tourette |

**Table S3:** list of all the edges of the network of co-alteration of opposing GM changes. Different nodes found in a same brain regions were labeled adding an increasing number.

| DECREASE<br>NODE<br>COORDINATES | DECREASE NODE LABEL | DECREASE NODE<br>RSN | INCREASE<br>NODE<br>COORDINATES | INCREASE NODE LABEL | INCREASE NODE<br>LABEL | PATEL'S K |
| --- | --- | --- | --- | --- | --- | --- |
| -2 -18 6 | Medial Dorsal Nucleus L | Thal/BG | 4 -44 56 | BA 5 1 R | DAN | 0.26 |
| -2 -18 6 | Medial Dorsal Nucleus L | Thal/BG | 10 2 2 | Medial Globus Pallidus R | Thal/BG | 0.55 |
| -2 -18 6 | Medial Dorsal Nucleus L | Thal/BG | -10 0 0 | Medial Globus Pallidus 2 L | Thal/BG | 0.43 |
| -2 -18 6 | Medial Dorsal Nucleus L | Thal/BG | 36 2 -20 | BA 38 R | S/VAN | 0.55 |
| -2 -18 6 | Medial Dorsal Nucleus L | Thal/BG | -26 -20 -20 | BA 35 L | Limbic | 0.43 |
| -2 -18 6 | Medial Dorsal Nucleus L | Thal/BG | -22 0 12 | Putamen 1 L | Thal/BG | 0.33 |
| -2 -18 6 | Medial Dorsal Nucleus L | Thal/BG | 0 40 -4 | BA 32 4 L | DMN | 0.43 |
| -2 -18 6 | Medial Dorsal Nucleus L | Thal/BG | -40 2 12 | BA 13 mid 2 L | S/VAN | 0.43 |
| -2 -18 6 | Medial Dorsal Nucleus L | Thal/BG | -4 -42 50 | BA 5 1 L | S/VAN | 0.33 |
| -2 -18 6 | Medial Dorsal Nucleus L | Thal/BG | -16 -50 58 | BA 7 L | DAN | 0.43 |
| -2 -18 6 | Medial Dorsal Nucleus L | Thal/BG | 16 10 4 | Putamen 1 R | Thal/BG | 0.43 |
| -2 -18 6 | Medial Dorsal Nucleus L | Thal/BG | 0 4 -2 | BA 25 L | Thal/BG | 0.55 |
| -2 -18 6 | Medial Dorsal Nucleus L | Thal/BG | -42 -18 -24 | BA 20 1 L | Limbic | 0.66 |
| -2 -18 6 | Medial Dorsal Nucleus L | Thal/BG | -28 -8 12 | BA 13 mid 5 L | Sensorimotor | 0.43 |
| -2 -18 6 | Medial Dorsal Nucleus L | Thal/BG | 22 2 2 | Putamen 2 R | Thal/BG | 0.33 |
| -2 -18 6 | Medial Dorsal Nucleus L | Thal/BG | -22 8 6 | Putamen 3 L | Thal/BG | 0.33 |
| -2 -18 6 | Medial Dorsal Nucleus L | Thal/BG | -34 -2 20 | BA 13 mid 4 L | S/VAN | 0.43 |
| -2 -18 6 | Medial Dorsal Nucleus L | Thal/BG | -16 34 -4 | BA 10 L | DMN | 0.43 |
| -42 16 0 | BA 13 ant 1 L | S/VAN | -18 -8 -14 | Amygdala L | Thal/BG | 0.49 |
| -42 16 0 | BA 13 ant 1 L | S/VAN | -28 -10 -2 | Putamen 4 L | Thal/BG | 0.61 |
| -42 16 0 | BA 13 ant 1 L | S/VAN | 10 2 2 | Medial Globus Pallidus R | Thal/BG | 0.24 |
| -42 16 0 | BA 13 ant 1 L | S/VAN | -26 -20 -20 | BA 35 L | Limbic | 0.34 |
| -42 16 0 | BA 13 ant 1 L | S/VAN | -22 0 12 | Putamen 1 L | Thal/BG | 0.49 |
| -42 16 0 | BA 13 ant 1 L | S/VAN | 40 -8 4 | BA 13 post 2 R | S/VAN | 0.49 |
| -42 16 0 | BA 13 ant 1 L | S/VAN | 30 -6 -2 | Putamen 1 R | S/VAN | 0.49 |
| -42 16 0 | BA 13 ant 1 L | S/VAN | -40 2 12 | BA 13 mid 2 L | S/VAN | 0.34 |
| -42 16 0 | BA 13 ant 1 L | S/VAN | 0 4 -2 | BA 25 L | Thal/BG | 0.49 |
| -42 16 0 | BA 13 ant 1 L | S/VAN | 4 -52 -16 | Culmen R | Cerebellum | 0.49 |
| -42 16 0 | BA 13 ant 1 L | S/VAN | -42 -18 -24 | BA 20 1 L | Limbic | 0.34 |
| -42 16 0 | BA 13 ant 1 L | S/VAN | -14 -16 -20 | BA 28 1 L | Limbic | 0.16 |
| -42 16 0 | BA 13 ant 1 L | S/VAN | -28 -8 12 | BA 13 mid 5 L | Sensorimotor | 0.61 |
| -42 16 0 | BA 13 ant 1 L | S/VAN | 26 -10 8 | Putamen 3 R | Thal/BG | 0.34 |
| -42 16 0 | BA 13 ant 1 L | S/VAN | -10 -10 0 | Thalamus L | Thal/BG | 0.49 |
| -42 16 0 | BA 13 ant 1 L | S/VAN | -22 8 6 | Putamen 3 L | Thal/BG | 0.24 |
| -42 16 0 | BA 13 ant 1 L | S/VAN | -28 26 14 | BA 13 ant 3 | S/VAN | 0.61 |
| -42 16 0 | BA 13 ant 1 L | S/VAN | -34 -2 20 | BA 13 mid 4 L | S/VAN | 0.61 |
| -42 16 0 | BA 13 ant 1 L | S/VAN | 20 0 -10 | BA 34 2 R | Limbic | 0.24 |
| -42 16 0 | BA 13 ant 1 L | S/VAN | -16 18 8 | Caudate Body 2 L | Thal/BG | 0.24 |
| -42 -8 8 | BA 13 mid 1 L | Sensorimotor | 40 -8 4 | BA 13 post 2 R | S/VAN | 0.57 |
| -42 -8 8 | BA 13 mid 1 L | Sensorimotor | 30 -6 -2 | Putamen 1 R | S/VAN | 0.57 |
| -42 -8 8 | BA 13 mid 1 L | Sensorimotor | 26 -10 8 | Putamen 3 R | Thal/BG | 0.45 |
| -42 -8 8 | BA 13 mid 1 L | Sensorimotor | 22 2 2 | Putamen 2 R | Thal/BG | 0.35 |
| -42 -8 8 | BA 13 mid 1 L | Sensorimotor | 20 0 -10 | BA 34 2 R | Limbic | 0.35 |
| -36 16 -8 | BA 47 1 L | DMN | 10 2 2 | Medial Globus Pallidus R | Thal/BG | 0.31 |
| -36 16 -8 | BA 47 1 L | DMN | -10 0 0 | Medial Globus Pallidus 2 L | Thal/BG | 0.41 |
| -36 16 -8 | BA 47 1 L | DMN | -26 -20 -20 | BA 35 L | Limbic | 0.41 |
| -36 16 -8 | BA 47 1 L | DMN | -22 0 12 | Putamen 1 L | Thal/BG | 0.54 |
| -36 16 -8 | BA 47 1 L | DMN | 40 -8 4 | BA 13 post 2 R | S/VAN | 0.54 |
| -36 16 -8 | BA 47 1 L | DMN | 30 -6 -2 | Putamen 1 R | S/VAN | 0.54 |
| -36 16 -8 | BA 47 1 L | DMN | -16 -50 58 | BA 7 L | DAN | 0.41 |
| -36 16 -8 | BA 47 1 L | DMN | 0 4 -2 | BA 25 L | Thal/BG | 0.54 |
| -36 16 -8 | BA 47 1 L | DMN | -28 -8 12 | BA 13 mid 5 L | Sensorimotor | 0.65 |
| -36 16 -8 | BA 47 1 L | DMN | 26 -10 8 | Putamen 3 R | Thal/BG | 0.41 |
| -36 16 -8 | BA 47 1 L | DMN | -12 -40 58 | BA 5 2 L | Sensorimotor | 0.41 |
| -36 16 -8 | BA 47 1 L | DMN | 22 2 2 | Putamen 2 R | Thal/BG | 0.31 |
| -36 16 -8 | BA 47 1 L | DMN | -22 8 6 | Putamen 3 L | Thal/BG | 0.31 |
| -36 16 -8 | BA 47 1 L | DMN | -28 26 14 | BA 13 ant 3 | S/VAN | 0.41 |
| -36 16 -8 | BA 47 1 L | DMN | -34 -2 20 | BA 13 mid 4 L | S/VAN | 0.41 |
| -36 16 -8 | BA 47 1 L | DMN | -16 18 8 | Caudate Body 2 L | Thal/BG | 0.31 |
| 46 10 -2 | BA 13 ant 1 R | S/VAN | 10 -42 66 | BA 5 2 R | Sensorimotor | 0.45 |
| -18 -8 -14 | Amygdala L | Thal/BG | 4 -44 56 | BA 5 1 R | DAN | 0.26 |
| -18 -8 -14 | Amygdala L | Thal/BG | 10 2 2 | Medial Globus Pallidus R | Thal/BG | 0.33 |
| -18 -8 -14 | Amygdala L | Thal/BG | -10 10 12 | Caudate Body L | Thal/BG | 0.66 |
| -18 -8 -14 | Amygdala L | Thal/BG | -22 0 12 | Putamen 1 L | Thal/BG | 0.33 |
| -18 -8 -14 | Amygdala L | Thal/BG | -4 -42 50 | BA 5 1 L | S/VAN | 0.33 |
| -18 -8 -14 | Amygdala L | Thal/BG | 16 10 4 | Putamen 1 R | Thal/BG | 0.43 |
| -18 -8 -14 | Amygdala L | Thal/BG | 6 -34 56 | BA 5 3 R | Sensorimotor | 0.55 |
| -18 -8 -14 | Amygdala L | Thal/BG | 22 2 2 | Putamen 2 R | Thal/BG | 0.33 |
| -18 -8 -14 | Amygdala L | Thal/BG | -22 8 6 | Putamen 3 L | Thal/BG | 0.33 |
| -18 -8 -14 | Amygdala L | Thal/BG | 20 0 -10 | BA 34 2 R | Limbic | 0.33 |
| -18 -8 -14 | Amygdala L | Thal/BG | 10 -42 66 | BA 5 2 R | Sensorimotor | 0.20 |
| -18 -8 -14 | Amygdala L | Thal/BG | -12 2 18 | Caudate Body 3 L | Thal/BG | 0.55 |
| 40 -56 38 | BA 40 R | DMN | -26 -20 -20 | BA 35 L | Limbic | 0.61 |
| 40 -56 38 | BA 40 R | DMN | -32 -12 -20 | Hippocampus L | Limbic | 0.61 |
| 40 -56 38 | BA 40 R | DMN | -22 22 0 | BA 47 5 L | ECN | 0.60 |
| 40 -56 38 | BA 40 R | DMN | -42 -18 -24 | BA 20 1 L | Limbic | 0.61 |
| 40 -56 38 | BA 40 R | DMN | -28 22 -10 | BA 47 3 L | DMN | 0.60 |
| 40 -56 38 | BA 40 R | DMN | -32 26 0 | BA 47 4 L | ECN | 0.61 |
| 40 -56 38 | BA 40 R | DMN | -16 34 -4 | BA 10 L | DMN | 0.61 |
| 40 -56 38 | BA 40 R | DMN | -16 18 8 | Caudate Body 2 L | Thal/BG | 0.60 |
| 2 38 30 | BA 9 1 R | DMN | -18 -8 -14 | Amygdala L | Thal/BG | 0.55 |
| 2 38 30 | BA 9 1 R | DMN | -28 -10 -2 | Putamen 4 L | Thal/BG | 0.43 |
| 2 38 30 | BA 9 1 R | DMN | 10 18 0 | Caudate Head R | Thal/BG | 0.43 |
| 2 38 30 | BA 9 1 R | DMN | -34 -52 46 | BA 40 L | DAN | 0.55 |
| 2 38 30 | BA 9 1 R | DMN | 14 -14 28 | BA 23 R | ECN | 0.55 |
| 0 18 42 | BA 6 L | ECN | -22 0 12 | Putamen 1 L | Thal/BG | 0.48 |
| 0 18 42 | BA 6 L | ECN | -28 -8 12 | BA 13 mid 5 L | Sensorimotor | 0.50 |
| 0 18 42 | BA 6 L | ECN | -22 8 6 | Putamen 3 L | Thal/BG | 0.48 |
| 0 18 42 | BA 6 L | ECN | -34 -2 20 | BA 13 mid 4 L | S/VAN | 0.50 |
| 0 18 42 | BA 6 L | ECN | -12 2 18 | Caudate Body 3 L | Thal/BG | 0.61 |
| 34 24 -6 | BA 47 1 R | ECN | 4 -44 56 | BA 5 1 R | DAN | 0.46 |
| 34 24 -6 | BA 47 1 R | ECN | -10 42 -4 | BA 32 5 L | DMN | 0.61 |
| 34 24 -6 | BA 47 1 R | ECN | -52 -24 -22 | BA 20 1 L | Limbic | 0.61 |
| 34 24 -6 | BA 47 1 R | ECN | 0 40 -4 | BA 32 4 L | DMN | 0.50 |
| 34 24 -6 | BA 47 1 R | ECN | 40 -8 4 | BA 13 post 2 R | S/VAN | 0.61 |
| 34 24 -6 | BA 47 1 R | ECN | -4 -42 50 | BA 5 1 L | S/VAN | 0.48 |
| 34 24 -6 | BA 47 1 R | ECN | 10 -42 66 | BA 5 2 R | Sensorimotor | 0.45 |
| -10 10 12 | Caudate Body 1 L | Thal/BG | 26 -10 8 | Putamen 3 R | Thal/BG | 0.61 |
| -10 10 12 | Caudate Body 1 L | Thal/BG | -28 26 14 | BA 13 ant 3 | S/VAN | 0.61 |
| -26 -20 -20 | BA 35 L | Limbic | -10 10 12 | Caudate Body L | Thal/BG | 0.48 |
| -26 -20 -20 | BA 35 L | Limbic | -22 0 12 | Putamen 1 L | Thal/BG | 0.39 |
| -26 -20 -20 | BA 35 L | Limbic | -4 -42 50 | BA 5 1 L | S/VAN | 0.39 |
| -26 -20 -20 | BA 35 L | Limbic | -14 -16 -20 | BA 28 1 L | Limbic | 0.37 |
| -26 -20 -20 | BA 35 L | Limbic | -22 8 6 | Putamen 3 L | Thal/BG | 0.39 |
| -26 -20 -20 | BA 35 L | Limbic | 20 0 -10 | BA 34 2 R | Limbic | 0.39 |
| -26 -20 -20 | BA 35 L | Limbic | -16 18 8 | Caudate Body 2 L | Thal/BG | 0.39 |
| -44 20 10 | BA 45 1 L | DMN | -18 -8 -14 | Amygdala L | Thal/BG | 0.58 |
| -44 20 10 | BA 45 1 L | DMN | -28 -10 -2 | Putamen 4 L | Thal/BG | 0.46 |
| -44 20 10 | BA 45 1 L | DMN | 4 -52 -16 | Culmen R | Cerebellum | 0.58 |
| -44 20 10 | BA 45 1 L | DMN | -28 26 14 | BA 13 ant 3 | S/VAN | 0.46 |
| -44 20 10 | BA 45 1 L | DMN | 10 -42 66 | BA 5 2 R | Sensorimotor | 0.28 |
| -16 34 24 | BA 32 1 L | DMN | -26 -20 -20 | BA 35 L | Limbic | 0.46 |
| -16 34 24 | BA 32 1 L | DMN | -32 -12 -20 | Hippocampus L | Limbic | 0.46 |
| -16 34 24 | BA 32 1 L | DMN | -22 22 0 | BA 47 5 L | ECN | 0.37 |
| -16 34 24 | BA 32 1 L | DMN | -42 -18 -24 | BA 20 1 L | Limbic | 0.46 |
| -16 34 24 | BA 32 1 L | DMN | -28 22 -10 | BA 47 3 L | DMN | 0.58 |
| -16 34 24 | BA 32 1 L | DMN | -32 26 0 | BA 47 4 L | ECN | 0.69 |
| -16 34 24 | BA 32 1 L | DMN | -34 -52 46 | BA 40 L | DAN | 0.58 |
| -16 34 24 | BA 32 1 L | DMN | -16 34 -4 | BA 10 L | DMN | 0.46 |
| -16 34 24 | BA 32 1 L | DMN | -16 18 8 | Caudate Body 2 L | Thal/BG | 0.37 |
| -16 34 24 | BA 32 1 L | DMN | 14 -14 28 | BA 23 R | ECN | 0.58 |
| 0 28 32 | BA 32 2 L | ECN | 10 18 0 | Caudate Head R | Thal/BG | 0.46 |
| 0 28 32 | BA 32 2 L | ECN | -34 -52 46 | BA 40 L | DAN | 0.58 |
| -2 30 -16 | BA 11 L | Limbic | -10 0 0 | Medial Globus Pallidus 2 L | Thal/BG | 0.48 |
| -2 30 -16 | BA 11 L | Limbic | -6 24 2 | BA 24 L | DMN | 0.60 |
| -60 -46 -4 | BA 21 L | ECN | 14 -10 50 | BA 6 2 R | Sensorimotor | 0.62 |
| -60 -46 -4 | BA 21 L | ECN | 4 -52 -16 | Culmen R | Cerebellum | 0.62 |
| -60 -46 -4 | BA 21 L | ECN | 20 0 -10 | BA 34 2 R | Limbic | 0.60 |
| 10 18 0 | Caudate Head R | Thal/BG | -22 0 12 | Putamen 1 L | Thal/BG | 0.60 |
| 11 18 0 | Caudate Head R | Thal/BG | -28 -8 12 | BA 13 mid 5 L | Sensorimotor | 0.61 |
| 12 18 0 | Caudate Head R | Thal/BG | -22 8 6 | Putamen 3 L | Thal/BG | 0.60 |
| 30 -6 -2 | Putamen 1 R | S/VAN | 28 -90 -6 | BA 18 R | Vis | 0.62 |
| -40 2 12 | BA 13 mid 2 L | S/VAN | -22 0 12 | Putamen 1 L | Thal/BG | 0.33 |
| -40 2 12 | BA 13 mid 2 L | S/VAN | 10 18 0 | Caudate Head R | Thal/BG | 0.43 |
| -40 2 12 | BA 13 mid 2 L | S/VAN | 16 10 4 | Putamen 1 R | Thal/BG | 0.43 |
| -40 2 12 | BA 13 mid 2 L | S/VAN | -22 22 0 | BA 47 5 L | ECN | 0.33 |
| -40 2 12 | BA 13 mid 2 L | S/VAN | -28 -8 12 | BA 13 mid 5 L | Sensorimotor | 0.43 |
| -40 2 12 | BA 13 mid 2 L | S/VAN | 20 12 -6 | Putamen 4 R | Limbic | 0.43 |
| -40 2 12 | BA 13 mid 2 L | S/VAN | 22 2 2 | Putamen 2 R | Thal/BG | 0.33 |
| -40 2 12 | BA 13 mid 2 L | S/VAN | -22 8 6 | Putamen 3 L | Thal/BG | 0.33 |
| -40 2 12 | BA 13 mid 2 L | S/VAN | -34 -2 20 | BA 13 mid 4 L | S/VAN | 0.43 |
| -40 2 12 | BA 13 mid 2 L | S/VAN | -12 2 18 | Caudate Body 3 L | Thal/BG | 0.55 |
| 20 -2 -24 | BA 28 R | Limbic | -4 -42 50 | BA 5 1 L | S/VAN | 0.48 |
| 32 4 0 | Clastrum 1 R | S/VAN | 28 -90 -6 | BA 18 R | Vis | 0.58 |
| 32 4 0 | Clastrum 1 R | S/VAN | 10 18 0 | Caudate Head R | Thal/BG | 0.46 |
| -34 12 6 | Clastrum L | S/VAN | 10 2 2 | Medial Globus Pallidus R | Thal/BG | 0.37 |
| -34 12 6 | Clastrum L | S/VAN | 16 10 4 | Putamen 1 R | Thal/BG | 0.46 |
| -34 12 6 | Clastrum L | S/VAN | 22 2 2 | Putamen 2 R | Thal/BG | 0.37 |
| -34 12 6 | Clastrum L | S/VAN | 10 -42 66 | BA 5 2 R | Sensorimotor | 0.28 |
| -48 6 32 | BA 9 1 L | DAN | -14 -16 -20 | BA 28 1 L | Limbic | 0.46 |
| -48 6 32 | BA 9 1 L | DAN | 10 -42 66 | BA 5 2 R | Sensorimotor | 0.45 |
| 30 14 0 | Clastrum 2 R | ECN | 10 2 2 | Medial Globus Pallidus R | Thal/BG | 0.60 |
| 30 14 0 | Clastrum 2 R | ECN | 16 10 4 | Putamen 1 R | Thal/BG | 0.61 |
| 30 14 0 | Clastrum 2 R | ECN | 22 2 2 | Putamen 2 R | Thal/BG | 0.60 |
| -32 -12 -20 | Hippocampus L | Limbic | -14 -16 -20 | BA 28 1 L | Limbic | 0.58 |
| -32 -12 -20 | Hippocampus L | Limbic | -28 22 -10 | BA 47 3 L | DMN | 0.39 |
| -32 -12 -20 | Hippocampus L | Limbic | -22 8 6 | Putamen 3 L | Thal/BG | 0.39 |
| -32 -12 -20 | Hippocampus L | Limbic | 20 0 -10 | BA 34 2 R | Limbic | 0.60 |
| -32 -12 -20 | Hippocampus L | Limbic | -16 18 8 | Caudate Body 2 L | Thal/BG | 0.39 |
| 0 4 -2 | BA 25 L | Thal/BG | -32 -12 -20 | Hippocampus L | Limbic | 0.50 |
| 4 -16 14 | Medial Dorsal Nucleus R | Thal/BG | 36 2 -20 | BA 38 R | S/VAN | 0.62 |
| 4 -16 14 | Medial Dorsal Nucleus R | Thal/BG | -26 -20 -20 | BA 35 L | Limbic | 0.61 |
| -2 42 44 | BA 8 1 L | DMN | 10 18 0 | Caudate Head R | Thal/BG | 0.50 |
| -2 42 44 | BA 8 1 L | DMN | -6 24 2 | BA 24 L | DMN | 0.61 |
| -2 42 44 | BA 8 1 L | DMN | 18 -60 -24 | Dentate | Cerebellum | 0.61 |
| -2 28 42 | BA 8 2 L | ECN | 10 2 2 | Medial Globus Pallidus R | Thal/BG | 0.35 |
| -2 28 42 | BA 8 2 L | ECN | -6 24 2 | BA 24 L | DMN | 0.57 |
| -2 28 42 | BA 8 2 L | ECN | 18 -60 -24 | Dentate | Cerebellum | 0.57 |
| -2 28 42 | BA 8 2 L | ECN | -34 -52 46 | BA 40 L | DAN | 0.57 |
| -2 28 42 | BA 8 2 L | ECN | 14 -14 28 | BA 23 R | ECN | 0.57 |
| 10 12 12 | Caudate Body R | Thal/BG | -22 0 12 | Putamen 1 L | Thal/BG | 0.48 |
| 10 12 12 | Caudate Body R | Thal/BG | -28 -8 12 | BA 13 mid 5 L | Sensorimotor | 0.50 |
| 10 12 12 | Caudate Body R | Thal/BG | 26 -10 8 | Putamen 3 R | Thal/BG | 0.50 |
| 10 12 12 | Caudate Body R | Thal/BG | -22 8 6 | Putamen 3 L | Thal/BG | 0.48 |
| 10 12 12 | Caudate Body R | Thal/BG | -28 26 14 | BA 13 ant 3 | S/VAN | 0.50 |
| 16 -8 -16 | BA 34 1 R | Limbic | -10 10 12 | Caudate Body L | Thal/BG | 0.48 |
| 16 -8 -16 | BA 34 1 R | Limbic | -22 8 6 | Putamen 3 L | Thal/BG | 0.39 |
| -44 10 18 | BA 13 ant 2 L | ECN | 14 -10 50 | BA 6 2 R | Sensorimotor | 0.61 |
| -44 10 18 | BA 13 ant 2 L | ECN | -14 -16 -20 | BA 28 1 L | Limbic | 0.46 |
| -44 10 18 | BA 13 ant 2 L | ECN | 10 -42 66 | BA 5 2 R | Sensorimotor | 0.45 |
| -6 40 24 | BA 9 2 L | DMN | -26 -20 -20 | BA 35 L | Limbic | 0.45 |
| -6 40 24 | BA 9 2 L | DMN | -20 14 -12 | BA 47 5 R | Limbic | 0.57 |
| -6 40 24 | BA 9 2 L | DMN | -32 -12 -20 | Hippocampus L | Limbic | 0.45 |
| -6 40 24 | BA 9 2 L | DMN | -22 22 0 | BA 47 5 L | ECN | 0.57 |
| -6 40 24 | BA 9 2 L | DMN | -42 -18 -24 | BA 20 1 L | Limbic | 0.45 |
| -6 40 24 | BA 9 2 L | DMN | -28 22 -10 | BA 47 3 L | DMN | 0.57 |
| -6 40 24 | BA 9 2 L | DMN | -32 26 0 | BA 47 4 L | ECN | 0.45 |
| -6 40 24 | BA 9 2 L | DMN | -34 -52 46 | BA 40 L | DAN | 0.57 |
| -6 40 24 | BA 9 2 L | DMN | -16 34 -4 | BA 10 L | DMN | 0.45 |
| -6 40 24 | BA 9 2 L | DMN | -16 18 8 | Caudate Body 2 L | Thal/BG | 0.35 |
| 48 14 8 | BA 44 R | S/VAN | -10 0 0 | Medial Globus Pallidus 2 L | Thal/BG | 0.61 |
| 48 14 8 | BA 44 R | S/VAN | -42 -18 -24 | BA 20 1 L | Limbic | 0.61 |
| 48 14 8 | BA 44 R | S/VAN | -6 24 2 | BA 24 L | DMN | 0.62 |
| -28 -82 4 | BA 18 L | Vis | 4 -44 56 | BA 5 1 R | DAN | 0.46 |
| -28 -82 4 | BA 18 L | Vis | -4 -42 50 | BA 5 1 L | S/VAN | 0.48 |
| -28 -82 4 | BA 18 L | Vis | 12 -50 54 | BA 7 L | DAN | 0.50 |
| -28 -82 4 | BA 18 L | Vis | -12 -40 58 | BA 5 2 L | Sensorimotor | 0.50 |
| -28 -82 4 | BA 13 post 1 L | Sensorimotor | 16 10 4 | Putamen 1 R | Thal/BG | 0.48 |
| -28 -82 4 | BA 13 post 1 L | Sensorimotor | 22 2 2 | Putamen 2 R | Thal/BG | 0.39 |
| 18 0 -34 | BA 38 R | Limbic | -4 -42 50 | BA 5 1 L | S/VAN | 0.48 |
| -14 -16 -20 | BA 28 1 L | Limbic | 4 -44 56 | BA 5 1 R | DAN | 0.46 |
| -14 -16 -20 | BA 28 1 L | Limbic | -10 10 12 | Caudate Body L | Thal/BG | 0.50 |
| -14 -16 -20 | BA 28 1 L | Limbic | -4 -42 50 | BA 5 1 L | S/VAN | 0.48 |
| -14 -16 -20 | BA 28 1 L | Limbic | 6 -34 56 | BA 5 3 R | Sensorimotor | 0.61 |
| -14 -16 -20 | BA 28 1 L | Limbic | 20 0 -10 | BA 34 2 R | Limbic | 0.48 |
| -14 -16 -20 | BA 28 1 L | Limbic | 10 -42 66 | BA 5 2 R | Sensorimotor | 0.45 |
| -8 -22 -2 | Pulvinar L | Thal/BG | 4 -44 56 | BA 5 1 R | DAN | 0.69 |
| -8 -22 -2 | Pulvinar L | Thal/BG | -10 42 -4 | BA 32 5 L | DMN | 0.61 |
| -8 -22 -2 | Pulvinar L | Thal/BG | -22 0 12 | Putamen 1 L | Thal/BG | 0.48 |
| -8 -22 -2 | Pulvinar L | Thal/BG | 0 40 -4 | BA 32 4 L | DMN | 0.50 |
| -8 -22 -2 | Pulvinar L | Thal/BG | -4 -42 50 | BA 5 1 L | S/VAN | 0.70 |
| -8 -22 -2 | Pulvinar L | Thal/BG | -42 -18 -24 | BA 20 1 L | Limbic | 0.50 |
| -8 -22 -2 | Pulvinar L | Thal/BG | 12 -50 54 | BA 7 L | DAN | 0.50 |
| -8 -22 -2 | Pulvinar L | Thal/BG | 6 -34 56 | BA 5 3 R | Sensorimotor | 0.61 |
| -8 -22 -2 | Pulvinar L | Thal/BG | 10 -42 66 | BA 5 2 R | Sensorimotor | 0.45 |
| -8 -22 -2 | Pulvinar L | Thal/BG | -12 2 18 | Caudate Body 3 L | Thal/BG | 0.61 |
| -18 -24 -8 | BA 28 2 L | DMN | 10 2 2 | Medial Globus Pallidus R | Thal/BG | 0.60 |
| -18 -24 -8 | BA 28 2 L | DMN | -10 10 12 | Caudate Body L | Thal/BG | 0.61 |
| -18 -24 -8 | BA 28 2 L | DMN | -4 -42 50 | BA 5 1 L | S/VAN | 0.60 |
| -18 -24 -8 | BA 28 2 L | DMN | 16 10 4 | Putamen 1 R | Thal/BG | 0.61 |
| -18 -24 -8 | BA 28 2 L | DMN | 22 2 2 | Putamen 2 R | Thal/BG | 0.60 |
| -40 -2 0 | BA 13 mid 3 L | S/VAN | 40 -8 4 | BA 13 post 2 R | S/VAN | 0.57 |
| -40 -2 0 | BA 13 mid 3 L | S/VAN | 30 -6 -2 | Putamen 1 R | S/VAN | 0.57 |
| -40 -2 0 | BA 13 mid 3 L | S/VAN | -22 22 0 | BA 47 5 L | ECN | 0.35 |
| -40 -2 0 | BA 13 mid 3 L | S/VAN | 26 -10 8 | Putamen 3 R | Thal/BG | 0.45 |
| -40 -2 0 | BA 13 mid 3 L | S/VAN | 22 2 2 | Putamen 2 R | Thal/BG | 0.35 |
| -40 -2 0 | BA 13 mid 3 L | S/VAN | 20 0 -10 | BA 34 2 R | Limbic | 0.35 |
| -34 18 -18 | BA 47 2 L | DMN | -16 -50 58 | BA 7 L | DAN | 0.50 |
| -34 18 -18 | BA 47 2 L | DMN | -12 -40 58 | BA 5 2 L | Sensorimotor | 0.50 |
| -34 18 -18 | BA 47 2 L | DMN | -10 -10 0 | Thalamus L | Thal/BG | 0.61 |
| -48 8 -2 | BA 22 2 L | S/VAN | 40 -8 4 | BA 13 post 2 R | S/VAN | 0.62 |
| -48 8 -2 | BA 22 2 L | S/VAN | 30 -6 -2 | Putamen 1 R | S/VAN | 0.62 |
| -48 8 -2 | BA 22 2 L | S/VAN | 26 -10 8 | Putamen 3 R | Thal/BG | 0.61 |
| -28 22 -10 | BA 47 3 L | DMN | -16 -50 58 | BA 7 L | DAN | 0.48 |
| -28 22 -10 | BA 47 3 L | DMN | 4 -52 -16 | Culmen R | Cerebellum | 0.60 |
| -28 22 -10 | BA 47 3 L | DMN | -12 -40 58 | BA 5 2 L | Sensorimotor | 0.48 |
| 0 -6 -6 | Hypothalamus L | Thal/BG | -32 -12 -20 | Hippocampus L | Limbic | 0.61 |
| 36 12 38 | BA 9 2 R | ECN | -54 -44 -12 | BA 20 2 L | ECN | 0.62 |
| -54 24 4 | BA 45 2 L | DMN | -28 26 14 | BA 13 ant 3 | S/VAN | 0.61 |
| -34 -72 0 | BA 19 L | Vis | 4 -44 56 | BA 5 1 R | DAN | 0.58 |
| -34 -72 0 | BA 19 L | Vis | 12 -50 54 | BA 7 L | DAN | 0.61 |
| 22 2 2 | Putamen 2 R | Thal/BG | 28 -90 -6 | BA 18 R | Vis | 0.61 |
| 44 28 28 | BA 9 2 R | ECN | -14 -16 -20 | BA 28 1 L | Limbic | 0.46 |
| -10 -10 0 | Thalamus L | Thal/BG | 4 -44 56 | BA 5 1 R | DAN | 0.58 |
| -10 -10 0 | Thalamus L | Thal/BG | -22 0 12 | Putamen 1 L | Thal/BG | 0.60 |
| -10 -10 0 | Thalamus L | Thal/BG | -4 -42 50 | BA 5 1 L | S/VAN | 0.60 |
| -10 -10 0 | Thalamus L | Thal/BG | 12 -50 54 | BA 7 L | DAN | 0.61 |
| -10 -10 0 | Thalamus L | Thal/BG | 10 -42 66 | BA 5 2 R | Sensorimotor | 0.57 |
| -10 -10 0 | Thalamus L | Thal/BG | -12 2 18 | Caudate Body 3 L | Thal/BG | 0.62 |
| -32 26 0 | BA 47 4 L | ECN | -28 -10 -2 | Putamen 4 L | Thal/BG | 0.50 |
| -32 26 0 | BA 47 4 L | ECN | -10 0 0 | Medial Globus Pallidus 2 L | Thal/BG | 0.50 |
| -32 26 0 | BA 47 4 L | ECN | -22 0 12 | Putamen 1 L | Thal/BG | 0.48 |
| -32 26 0 | BA 47 4 L | ECN | -28 -8 12 | BA 13 mid 5 L | Sensorimotor | 0.50 |
| -32 26 0 | BA 47 4 L | ECN | -34 -2 20 | BA 13 mid 4 L | S/VAN | 0.50 |
| -16 -26 -18 | Culmen L | Cerebellum | -10 10 12 | Caudate Body L | Thal/BG | 0.61 |
| -16 -26 -18 | Culmen L | Cerebellum | -4 -42 50 | BA 5 1 L | S/VAN | 0.60 |
| -16 -26 -18 | Culmen L | Cerebellum | 20 0 -10 | BA 34 2 R | Limbic | 0.60 |
| 8 32 24 | BA 32 1 R | ECN | -26 -20 -20 | BA 35 L | Limbic | 0.46 |
| 8 32 24 | BA 32 1 R | ECN | -22 0 12 | Putamen 1 L | Thal/BG | 0.37 |
| 8 32 24 | BA 32 1 R | ECN | -14 -16 -20 | BA 28 1 L | Limbic | 0.31 |
| 8 32 24 | BA 32 1 R | ECN | -28 -8 12 | BA 13 mid 5 L | Sensorimotor | 0.46 |
| 8 32 24 | BA 32 1 R | ECN | -22 8 6 | Putamen 3 L | Thal/BG | 0.37 |
| -18 -14 -6 | Medial Globus Pallidus 1 L | Thal/BG | 4 -44 56 | BA 5 1 R | DAN | 0.58 |
| -18 -14 -6 | Medial Globus Pallidus 1 L | Thal/BG | 10 2 2 | Medial Globus Pallidus R | Thal/BG | 0.60 |
| -18 -14 -6 | Medial Globus Pallidus 1 L | Thal/BG | -10 10 12 | Caudate Body L | Thal/BG | 0.61 |
| -18 -14 -6 | Medial Globus Pallidus 1 | Thal/BG | 16 10 4 | Putamen 1 R | Thal/BG | 0.61 |
|  | L |  |  |  |  |  |
| -18 -14 -6 | Medial Globus Pallidus 1<br>L | Thal/BG | 6 -34 56 | BA 5 3 R | Sensorimotor | 0.62 |
| -18 -14 -6 | Medial Globus Pallidus 1<br>L | Thal/BG | 22 2 2 | Putamen 2 R | Thal/BG | 0.60 |
| -18 -14 -6 | Medial Globus Pallidus 1<br>L | Thal/BG | 10 -42 66 | BA 5 2 R | Sensorimotor | 0.57 |
| -40 26 -6 | BA 47 4 L | DMN | -28 -10 -2 | Putamen 4 L | Thal/BG | 0.45 |
| -40 26 -6 | BA 47 4 L | DMN | -10 0 0 | Medial Globus Pallidus 2 L | Thal/BG | 0.68 |
| -40 26 -6 | BA 47 4 L | DMN | -22 0 12 | Putamen 1 L | Thal/BG | 0.57 |
| -40 26 -6 | BA 47 4 L | DMN | 40 -8 4 | BA 13 post 2 R | S/VAN | 0.57 |
| -40 26 -6 | BA 47 4 L | DMN | 30 -6 -2 | Putamen 1 R | S/VAN | 0.57 |
| -40 26 -6 | BA 47 4 L | DMN | -40 2 12 | BA 13 mid 2 L | S/VAN | 0.45 |
| -40 26 -6 | BA 47 4 L | DMN | 0 4 -2 | BA 25 L | Thal/BG | 0.57 |
| -40 26 -6 | BA 47 4 L | DMN | -28 -8 12 | BA 13 mid 5 L | Sensorimotor | 0.68 |
| -40 26 -6 | BA 47 4 L | DMN | 26 -10 8 | Putamen 3 R | Thal/BG | 0.45 |
| -40 26 -6 | BA 47 4 L | DMN | -10 -10 0 | Thalamus L | Thal/BG | 0.57 |
| -40 26 -6 | BA 47 4 L | DMN | -22 8 6 | Putamen 3 L | Thal/BG | 0.35 |
| -40 26 -6 | BA 47 4 L | DMN | -28 26 14 | BA 13 ant 3 | S/VAN | 0.45 |
| -40 26 -6 | BA 47 4 L | DMN | -34 -2 20 | BA 13 mid 4 L | S/VAN | 0.68 |
| -40 26 -6 | BA 47 4 L | DMN | -16 18 8 | Caudate Body 2 L | Thal/BG | 0.35 |
| 10 30 34 | BA 6 1 R | ECN | -14 -16 -20 | BA 28 1 L | Limbic | 0.58 |
| -12 36 14 | BA 32 3 L | DMN | -26 -20 -20 | BA 35 L | Limbic | 0.61 |
| -12 36 14 | BA 32 3 L | DMN | -32 -12 -20 | Hippocampus L | Limbic | 0.61 |
| -12 36 14 | BA 32 3 L | DMN | -22 22 0 | BA 47 5 L | ECN | 0.60 |
| -12 36 14 | BA 32 3 L | DMN | -42 -18 -24 | BA 20 1 L | Limbic | 0.61 |
| -12 36 14 | BA 32 3 L | DMN | -28 22 -10 | BA 47 3 L | DMN | 0.60 |
| -12 36 14 | BA 32 3 L | DMN | -32 26 0 | BA 47 4 L | ECN | 0.61 |
| -12 36 14 | BA 32 3 L | DMN | -16 34 -4 | BA 10 L | DMN | 0.61 |
| -12 36 14 | BA 32 3 L | DMN | -16 18 8 | Caudate Body 2 L | Thal/BG | 0.60 |

**Table S4:** list of medications taken by the subjects of the experiments that were included or excluded in the meta-analysis of the co-alteration between opposing GM changes.

| Included | Author | Diagnosis | Medicated patients/ total | Pharmacological treatment (number of subjects) |
| --- | --- | --- | --- | --- |
| yes | BrieberS, 2007 | Attention Deficit/Hyperactivity Disorder | 10 on 15 | Risperidone (2), Psychostimulants (Methylphenidate, 10) |
| yes | WangJ, 2007 | Attention Deficit/Hyperactivity Disorder | 0 on 12 | none |
| yes | AdlerCM, 2005 | Bipolar Disorder | 23 on 32 | Lithium, Divalproex, Stabilizers, anti-seizures, antidepressants, benzodiazepines, atypical antipsychotics |
| yes | CuiL, 2011 | Bipolar Disorder | 0 on 24 | Stabilizers (lithium, sodium valproate, 17), antipsychotics (Quetiapine, Olanzapine Clozapine, 10), but note at the time of the scanning |
| yes | HaldaneM, 2008 | Bipolar Disorder | 44 on 44 | Antipsychotics (Chlorpromazine, 22), stabilizers (Lithium, Valproate, Carbamazepine, 31), antidepressants (8) |
| yes | LadouceurCD, 2008 | Bipolar Disorder | 0 on 20 | none |
| yes | SaricicekA, 2015 | Bipolar Disorder | Unspecified on 28 | Lithium (18), Valproate (12), Antipsychotics (14). Antidepressant (2) |
| yes | TangLR, 2014 | Bipolar Disorder | 27 on 27 | Antidepressants (Citalopram, 7; Sertraline, 8; Paroxetine, 1), Mood-stabilizer (lithium, 9; Sodium valproate, 6; Divalproex, 15), Antipsychotics (Quetiapine, 10; Olanzapine, 1; Risperidone, 5) |
| yes | WatsonDR, 2012 | Bipolar Disorder | 21 on 24 | Atypical antipsychotics (Risperidone, 6; Olanzapine, 10; Quetiapine, 3), Lithium (2) |
| yes | ArnoneD, 2009 | Major Depressive Disorder | 0 on 25 | none |
| yes | ArnoneD, 2013 | Major Depressive Disorder | 0 on 39 | none |
| yes | GongQ, 2011 | Major Depressive Disorder | 23 on 23 | Antidepressants (Tricyclic, SNRI, SRI) |
| yes | GongQ, 2011 | Major Depressive Disorder | 23 on 23 | Antidepressants (Tricyclic, SNRI, SRI) |
| yes | HwangJ, 2010 | Major Depressive Disorder | Unspecified on 26 | none reported |
| yes | LeungKK, 2009 | Major Depressive Disorder | 17 on 17 | SSRIs, tricyclic antidepressants, antipsychotics, benzodiazepine or hypnotics, or a combination of these |
| yes | ScheuereckerJ, 2010 | Major Depressive Disorder | 0 on 13 | none |
| yes | GilbertAR, 2008 | Obsessive Compulsive Disorder | 20 on 20 | Antidepressants (20: Fluoxetine, 9; Paroxetine, 4; Clomipramine, 2; Sertraline, Citalopram, Amitriptyline, 1), Anxiolytics (Diazepam, 2; Tamazepam, 1; Diazepam & Zopiclone, 1), Lithium + Sertraline (1), Thyroxine + Fluoxetine (1) |
| yes | PujolJ, 2004 | Obsessive Compulsive Disorder | Unspecified on 72 | Yes, but unspecified |
| yes | SzeszkoPR, 2008 | Obsessive Compulsive Disorder | 0 on 26 | none |
| yes | ValenteAAJr, | Obsessive | 11 on 19 | SSRI (4), Clomipramine (7) |
|  | 2005 | Compulsive Disorder |  |  |
| yes | ValenteAAJr, 2005 | Obsessive Compulsive Disorder | 11 on 19 | SSRI (4), Clomipramine (7) |
| yes | YooSY, 2008 | Obsessive Compulsive Disorder | 0 on 47 | Previously: Fluoxetine (19), Paroxetine (7), Fluvoxamine (6), Sertraline(27) |
| yes | LudolphAG, 2006 | Other Disorders of Psychological Development. Tourette | 4 on 14 | Stimulants (4) |
| yes | MarcelisM, 2003 | Other psychotic disorder not due to a substance or known physiological condition | 27 on 27 | Antipsychotics (13 Typical, 13 atypical, 1 mixed) |
| yes | AbellF, 1999 | Pervasive Developmental Disorders, Autism | 0 on 15 | none |
| yes | BrieberS, 2007 | Pervasive Developmental Disorders, Autism | 2 on 15 | Risperidone (2) |
| yes | ChengY, 2011 | Pervasive Developmental Disorders, Autism | 0 on 25 | none |
| yes | ChengY, 2011 | Pervasive Developmental Disorders, Autism | 0 on 11 | none |
| yes | ChengY, 2011 | Pervasive Developmental Disorders, Autism | 0 on 12 | none |
| yes | ChengY, 2011 | Pervasive Developmental Disorders, Autism | 0 on 12 | none |
| yes | EckerC, 2010 | Pervasive Developmental Disorders, Autism | 0 on 22 | none |
| yes | EckerC, 2012 | Pervasive Developmental Disorders, Autism | 0 on 89 | none |
| yes | HydeKL, 2010 | Pervasive Developmental Disorders, Autism | 0 on 13 | none |
| yes | KeX, 2008 | Pervasive Developmental Disorders, Autism | 0 on 15 | none |
| yes | MengottiP, 2011 | Pervasive Developmental Disorders, Autism | 0 on 20 | none (sedation when necessary in the MR) |
| yes | SalmondCH, 2007 | Pervasive Developmental Disorders, Autism | 0 on 9 | none |
| yes | ToalF, 2010 | Pervasive Developmental Disorders, Autism | 0 on 26 | none |
| yes | AntonovaE, 2005 | Schizophrenia | 40 on 40 | Antipsychotics (Chlorpromazine, 40), anti-cholinergic (Procyclidine, 15) |
| yes | BassittDP, 2007 | Schizophrenia | 30 on 30 | Typical antipsychotics (4), second-generation antipsychotics (17), Clozapine (21), typical plus atypical antipsychotics (6), typical antipsychotic plus Clozapine (2) |
| yes | CuiL, 2011 | Schizophrenia | 23 on 23 | Risperidone, Quetiapine, Olanzapine, Sulpiride, Clozapine |
| yes | DengMY, 2009 | Schizophrenia | 10 on 10 | Typical (Flupenthixol, Trifluoperazine, Haloperidol 6) and atypical (5) antipsychotics, Amisulpride (11) |
| yes | DengMY, 2009 | Schizophrenia | 10 on 10 | Typical (Flupenthixol, Trifluoperazine, Haloperidol 6) and atypical (5) antipsychotics, Amisulpride (11) |
| yes | GiulianiNR, 2005 | Schizophrenia | Unspecified on 34 | none reported |
| yes | HaTH, 2004 | Schizophrenia | 33 on 35 | Risperidone (21), Olanzapine (9), Chlozapine (3) |
| yes | HoneaRA, 2008 | Schizophrenia | 167 on 169 | Antipsychotics (typical, 22; atypical, 136; mixed, 9) |
| yes | HulshoffPolHE, 2001 | Schizophrenia | 155 on 158 | Antipsychotics (Clozapine, Risperidone, Olanzapine, Sertindole) |
| yes | KasperekT, 2010 | Schizophrenia | 49 on 49 | Antipsychotics (all, Risperidone, Olanzapine), benzodiazepines, hypnotics, anticholinergic antiparkinsonics |
| yes | KawasakiY, 2004 | Schizophrenia | 25 on 25 | Antipsychotics (9 atypical, 14 atypical) |
| yes | MolinaV, 2011 | Schizophrenia | 24 on 24 | Haloperidol (26), Pimocide (4), Levomepromazine (3), Thioridazine (2), Risperidone (3) |
| yes | MolinaV, 2011 | Schizophrenia | 30 on 30 | Haloperidol (26), Pimocide (4), Levomepromazine (3), Thioridazine (2), Risperidone (3) |
| yes | O'DalyO, 2007 | Schizophrenia | 26 on 28 | Atypical antipsychotics (21), typical antipsychotics, (5) |
| yes | PriceG, 2010 | Schizophrenia | 47 on 47 | Antipsychotics (all, Olanzapine, 23; Risperidone, 16; Aripiprazole, 4; Amisulpride, 3; Quetiapine, 1; Haloperidol, 1), antidepressants (7) |
| yes | Salgado-PinedaP, 2003 | Schizophrenia | 0 on 13 | none |
| yes | SchifferB, 2013 | Schizophrenia | 24 on 25 | Antipsychotics |
| yes | ShapleskeJ, 2002 | Schizophrenia | Unspecified on 31 | none reported |
| yes | ShapleskeJ, 2002 | Schizophrenia | Unspecified on 32 | none reported |
| yes | SmesnyS, 2010 | Schizophrenia | 0 on 13 | previously: Risperidone (3), unknown (1) |
| yes | SmesnyS, 2010 | Schizophrenia | 0 on 11 | previously: Risperidone (2), Olanzapine (4) , Haloperidol (1), Perazine (1), unknown (1) |
| yes | SuzukiM, 2002 | Schizophrenia | 42 on 42 | Haloperidol or equivalent (34), Risperidone (19), Quetiapine (1) |
| yes | WatsonDR, 2012 | Schizophrenia | 25 on 25 | Typical (1), atypical antipsychotics (24) |
| yes | WhitfordTJ, 2006 | Schizophrenia | 41 on 41 | Amisulpride, Quetiapine, Olanzapine, Risperidone, Clozapine |
| yes | WilkeM, 2001 | Schizophrenia | 43 on 48 | Atypical (22), typical (16), combinations (5), 13 of the atypical or no medication had typical neuroleptics in the past |
| yes | LuC, 2010 | Specific Developmental Disorders of Speech and Language | 0 on 12 | none |
| yes | WatkinsKE, 2002 | Specific Developmental Disorders of Speech and Language | 0 on 10 | none |
| no | GranertO, 2011 | Dystonia | 0 on 11 | previously: Botulyn (4) |
| no | ObermannM, 2007 | Dystonia | 17 on 20 | Botulinum toxin A |
| no | ChanCH, 2006 | Epilepsy and Recurrent Seizures | 12 on 13 | Antiepileptics (12) |
| no | deAraujo-FilhoGM, 2009 | Epilepsy and Recurrent Seizures | 16 on 16 | Adequate antiseizure drugs (Valproate, Topiramate, Lamotrigine, Clonazepam, 15), inadequate pharmacological treatment (1) |
| no | KellerSS, 2002 | Epilepsy and Recurrent Seizures | Unspecified on 40 | non controlled for |
| no | KellerSS, 2002 | Epilepsy and Recurrent Seizures | Unspecified on 36 | non controlled for |
| no | KellerSS, 2002 | Epilepsy and Recurrent Seizures | Unspecified on 58 | non controlled for |
| no | KellerSS, 2002 | Epilepsy and Recurrent Seizures | Unspecified on 58 | non controlled for |
| no | KellerSS, 2002 | Epilepsy and Recurrent Seizures | Unspecified on 58 | non controlled for |
| no | KimJH, 2007 | Epilepsy and Recurrent Seizures | 25 on 25 | Antiepileptics (Valproate, 17; Lamotrigine, 6) |
| no | LinK, 2009 | Epilepsy and Recurrent Seizures | 30 on 30 | Antiepileptics (Valproate, Lamotrigine, Carbamazepine, Topiramate, Clobazam, Clonazepam, Phenytoin, Phenobarbital) |
| no | LinK, 2009 | Epilepsy and Recurrent Seizures | 19 on 19 | Antiepileptics (Valproate, Lamotrigine, Carbamazepine, Topiramate, Clobazam, Clonazepam, Phenytoin, Phenobarbital) |
| no | LinK, 2009 | Epilepsy and Recurrent Seizures | 30 on 30 | Antiepileptics (Valproate, Lamotrigine, Carbamazepine, Topiramate, Clobazam, Clonazepam, Phenytoin, Phenobarbital) |
| no | RiedererF, 2008 | Epilepsy and Recurrent Seizures | Unspecified on 12 | none reported |
| no | RiedererF, 2008 | Epilepsy and Recurrent Seizures | Unspecified on 12 | none reported |
| no | CelleS, 2010 | Other<br>extrapyramidal<br>and movement<br>disorders | 0 on 17 | none |
| no | EtgenT, 2005 | Other<br>extrapyramidal<br>and movement<br>disorders | Unspecified<br>on 28 | L-dopa, dopamine agonists or opioids |
| no | LinCH, 2013 | Other<br>extrapyramidal<br>and movement<br>disorders | Unspecified<br>on 10 | none reported |
| no | LinCH, 2013 | Other<br>extrapyramidal<br>and movement<br>disorders | Unspecified<br>on 10 | none reported |
| no | LinCH, 2013 | Parkinson's<br>Disease | Unspecified<br>on 10 | previously: L-dopa (number unspecified) |
| no | LinCH, 2013 | Parkinson's<br>Disease | Unspecified<br>on 10 | none reported |
